# Unmasking Supervillin*: SVIL* haploinsufficiency causes hypertrophic cardiomyopathy by impairing mechanotransduction and cellular energetics

**DOI:** 10.64898/2026.07.01.735949

**Authors:** Yifan J. Li, Yiangos Psaras, Violetta Steeples, Josephine M. Watkins, Charlotte Hooper, Marta Moya-Jódar, Thomas Nicol, Alexander J. Sparrow, Marcos Garcia-Lacarte, Samuel TM Jones, Isabelle Bond, Niklas Beyhoff, Paul Robinson, Marieluise Kirchner, Philipp Mertins, James S Ware, R Thomas Lumbers, Betty Raman, Hugh Watkins, Christopher N. Toepfer

## Abstract

**Background:** Rare heterozygous loss-of-function (LoF) variants in *SVIL*, encoding the Z-disk and costameric protein supervillin, have recently been identified as a cause of hypertrophic cardiomyopathy (HCM). Although supervillin is implicated in actin-dependent mechanotransduction, the mechanisms linking *SVIL* deficiency to cardiomyopathy remain poorly understood. Homozygous LoF cause a novel skeletal Myofibrillar Myopathy-10 (MFM-10) while heterozygous LoF cause HCM without skeletal myopathy. In this study we use a human model system to disentangle the LoF pathomechanism of the scaffolding protein supervillin in cardiomyocytes and its clinical implications.

**Methods:** Using CRISPR/Cas-9 we engineered a representative pathogenic LoF variant Q255X into an isogenic induced pluripotent stem cell (iPSC) line creating the heterozygous *SVIL*^Q255X/+^ and homozygous *SVIL*^Q255X/Q255X^ cell lines. These lines were differentiated into iPSC-derived cardiomyocytes (iPSC-CMs) and cellular phenotypes were assessed using bulk RNA-sequencing, LC-MS proteomics, electrophysiological and calcium handling analyses, contractility measurements, sarcomere organization analysis, Seahorse metabolic flux assay, and pharmacological intervention with mavacamten.

**Results:** The Q255X variant resulted in *SVIL* haploinsufficiency at both RNA and protein levels with no evidence of a truncated protein. Compared with isogenic controls, *SVIL*^Q255X/+^ iPSC-CMs demonstrated action potential shortening, calcium transient elongation, sarcomeric disorganization and hypertrophy, and impaired mitochondrial respiration. Multi-omic analyses of *SVIL*^Q255X/+^ iPSC-CMs showed a profile of cellular stress and inflammation, hypertrophic and pro-fibrotic signalling, and a pseudohypoxic state driven by decreased respiration and a HIF-induced glycolytic shift. These abnormalities were not present in *SVIL*^Q255X/Q255X^ cardiomyocytes, consistent with a relatively limited cardiac phenotype reported in homozygous variant carriers. Mavacamten improved sarcomeric disorganization and hypertrophy in *SVIL*^Q255X/+^ cells but did not rescue energetic compromise.

**Conclusions:** Pathogenic heterozygous *SVIL* LoF produces a distinct cellular phenotype characterized by impaired mechanotransduction, mitochondrial dysfunction, and maladaptive metabolic remodelling that promotes hypertrophic and pro-fibrotic signalling. These findings define a mechanistic basis for *SVIL*-associated cardiomyopathy and identify metabolic dysfunction as a potential therapeutic target beyond sarcomere-directed therapy.

**Clinical Perspective:** *What Is New?:*

- *SVIL* haploinsufficiency causes HCM through a mechanism distinct from canonical sarcomeric disease, characterized by impaired mechanotransduction, mitochondrial dysfunction, and pseudohypoxia-driven metabolic remodeling.
- Heterozygous *SVIL* loss of function produces a substantially more severe cardiomyocyte phenotype than homozygous loss of function, providing a mechanistic explanation for the predominance of cardiac disease in heterozygous variant carriers.
- Mavacamten improves sarcomeric organization but does not restore impaired mitochondrial respiration, demonstrating that energetic dysfunction persists despite sarcomere-directed therapy.

*What Are the Clinical Implications?:*

- Our findings give functional evidence to support *SVIL* as a clinically relevant HCM disease gene and its inclusion in clinical genetic testing panels.
- These findings establish *SVIL*-associated cardiomyopathy as a mechanistically distinct form of HCM and offer insight into the pathomechanism of Z-disk and costameric HCM
- The persistence of mitochondrial dysfunction despite myosin inhibition suggests that drugs targeting mitochondrial bioenergetics may be a therapeutic strategy in patients with *SVIL*-associated cardiomyopathy.

## Introduction

HCM is a complex genetic disease with both monogenic and polygenic drivers. Rare pathogenic variants in eight core sarcomeric genes cause most monogenic disease accounting for ∼35% of cases (HCM_SARC+_), and common variant contribution across non-sarcomeric genes cause polygenic disease (HCM_SARC-_). However, a subset of non-core sarcomeric HCM genes underlie both monogenic and polygenic disease^1,2^. Such genes are usually involved in sarcomere regulation, (e.g. *FHOD3*^3^*, ALPK3*^4^), calcium signalling or cellular stress response (e.g. *PLN, CRYAB*), or are Z-disc/costamere related genes (e.g. *CSRP3*, *FHL1, FLNC, ACTN2, MYOZ2*)^5,6^.

Recently, a meta-analysis of genome wide association studies (GWAS) with rare variant association identified *SVIL* (encoding the multi-domain, actin-binding Z-disc protein supervillin) as a novel HCM disease gene, with rare truncating variants in heterozygous patients conferring a 10.5-fold increased risk of HCM^2^. Located on the reverse strand of chromosome 10, *SVIL* has multiple protein-coding transcripts of which the most expressed in the heart is the skeletal and cardiac muscle-specific *SVIL-201* (Ensembl, release 114^7^) encoding a ∼250kDa protein (also called Archvillin^8^), followed by the smooth muscle and non-muscle-specific *SVIL-203* (∼200kDa). Bulk tissue gene expression for *SVIL* (GTEx Portal version 10^9^) shows the highest levels of *SVIL* expression in skeletal muscle, followed by smooth muscle (across a range of organs) and cardiac muscle (**Supplemental Figure 1**). Supervillin binds to filamentous (F)-actin, localises to the Z-disc and nucleus and co-localises with dystrophin and nebulin at the costamere, functioning as an additional linkage required for efficient lateral force transduction between actin filaments and the sarcolemma^8,10^. Further roles of supervillin in myocytes include myofibrillogenesis^11^, and in non-myocytes, cytokinesis^12,13^, cell adhesion^14^, spreading^15^, motility^16^, and survival^17^. A previous study using machine learning identified *SVIL* as a candidate gene affecting LV diameter, with morpholino knockdown producing cardiac oedema in zebrafish^18^. In humans, *SVIL* LoF is associated with a smaller descending aorta diameter^19^ and homozygous LoF observed in four patients displayed a novel skeletal myopathy termed Myofibrillar Myopathy-10 (MFM-10) with mild cardiac features^20^. The *SVIL* locus also reached GWAS significance for non-ischaemic cardiomyopathy independent of HCM in a recent study, with functional study using si-RNA knockdown demonstrating murine cardiomyocyte hypertrophy, apoptosis and impaired viability^21^.

Little is known about the mechanism by which *SVIL* LoF precipitates HCM. The previously described unifying mechanism of pathological hypertrophy in HCM is one of hypercontractility and diastolic insufficiency driving inefficient energy usage, leading to energy depletion and calcium dysregulation and converging on downstream hypertrophic signalling pathways^22^. This model fits to the more recent and classical pathomechanisms discovered that account for typical sarcomeric hypercontractile HCM phenotypes caused by missense *MHY7*^23^ or truncating *MYBPC3* variants^24,25^, and somewhat accounts for thin filament HCM variants that drive HCM via destabilisation of calcium regulation directly^26–28^. However, further mechanistic insight is required to explain how partial loss of a Z-disk and costameric actin interacting protein such as supervillin drives HCM disease processes that are not evident in full loss of function (LoF).

Here we use iPSC-CMs in combination with CRISPR/Cas-9 to model *SVIL*^Q255X/+^ and *SVIL*^Q255X/Q255X^ patient variants in the dish. In doing so we demonstrate for the first time the consequences of heterozygous and homozygous *SVIL* LoF at the RNA, protein, and cell function levels, with bulk transcriptomic and proteomic data suggesting *SVIL* cardiomyopathy to be associated with heterozygous variant carriers. Thus, we propose a novel mechanism of pathological cardiac hypertrophy in line with the energy depletion hypothesis that is unique and specific to heterozygous *SVIL* LoF and establish the mechanistic framework for therapeutic interventions.

## Methods

### Generation of isogenic iPSC-CM SVIL truncation model

The LoF *SVIL* variant (p.Gln255X) carried by two cousins with HCM described in Tadros et. al.^2^ was engineered into the HPSI0114i-kolf_2 (KOLF2) iPSC line using CRISPR/Cas-9 with homology directed repair (HDR) as previously described^29^, generating the isogenic wild-type (WT) *SVIL*^+/+^, *SVIL*^Q255X/+^ and *SVIL*^Q255X/Q255X^ lines (**See Supplemental Methods**). iPSCs were differentiated into cardiomyocytes (CMs) as previously described^30,31^ (**See Supplemental Methods**).

### Immunofluorescence analyses of isogenic iPSC-CMs

Replated iPSC-CMs were fixed in 4% paraformaldehyde, permeabilised and stained using anti-*SVIL* (Sigma-Aldrich, HPA020138-25UL), anti-ACTN2 (Sigma-Aldrich, EA53), or rhodamine phalloidin (Invitrogen™, R415) **(Supplemental Table 1).** Mouse and pig LV tissue were used as positive controls and pig liver tissue was used as negative control (**Supplemental Figure 2**). For mitochondrial visualisation and nuclear studies, fixed cells were stained with the mitochondrial marker ATP5A (Abcam, Ab14748), anti-sarcomeric alpha actinin (Abcam, ab68167), and Hoechst 33342 (ThermoFisher, R37605). Imaging was done at 63x using a Leica SP5 confocal microscope. Analysis was performed in Fiji (Image J, NIH; version 1.54p) by applying automated thresholding and segmentation and measurements done using built-in analysis tools.

### Functional analyses: action potential, calcium transients, contractility, and sarcomere organization

Replated iPSC-CMs were used for live-cell imaging between D30-33 (**See Supplemental Methods**). The FluoVolt™ Membrane Potential Kit ((ThermoFisher Scientific, F10488) was used for membrane action potential (AP) measurements. For calcium transients and 2D contractility, iPSC-CMs were transduced using replication deficient adenoviral constructs (AV) encoding RGECO (a red fluorescent calcium indicator) and α-actinin-GFP fusion protein as previously reported^27^. Acute drug treatment was performed using 1 μM mavacamten in dimethyl sulfoxide (DMSO) for 1 hour. A single video is defined as a technical replicate and an iPSC-CM differentiation batch is defined as a biological replicate. AP videos were analysed using AP Track (a modified version of Caltrack^27^), calcium using CalTrack^27^, contractility using SarcTrack^25^, and sarcomere organisation using Sarcomere Analysis Multitool (SarcAsM, version 0.2.0)^32^. Engineered heart tissues (EHTs) were cast as previously described^30^ with modifications (**See Supplemental Methods**) and analysed using BeatProfiler^33^.

### DRX and SRX analysis using MANT-ATP

The MANT ATP (fluorescent ATP analogue) assay was used to investigate the super-relaxed (SRX) and disordered-relaxed (DRX) state of myosin per published protocol^34^. Briefly, non-replated beating sheets of iPSC-CMs cultured in 6-well plates to D30-33 were dissected into flow chambers, skinned, and maintained in glycerinating solution. Rigor buffer, MANT-ATP (ThermoFisher Scientific, M12417) and chase (non-fluorescent) ATP solutions were used for fluorescence-decay imaging obtained using above-mentioned Nikon/Photometrics setup.

### Seahorse mitochondrial metabolism flux analysis

Cellular mitochondrial oxygen consumption rate (OCR) was assessed using Seahorse XF Mito Stress Test kit (Agilent, 103015-100) on the Seahorse XF Pro Analyzer per Agilent user guide. Passaged iPSC-CMs at D17 were seeded at 3000 cells/well density onto XFe96/XF Pro Cell Culture microplates and cultured until D25. 24-hour drug treatments using 1 µM mavacamten were performed as previously described^23^ (**See Supplemental Methods**).

### RNA studies

Replated iPSC-CMs cultured to D30-33 were processed for RNA extraction and qPCR was performed using TaqMan™ probes for *SVIL* and β-tubulin as control (**See Supplemental Method**). Extracted RNA samples were sent for lncRNA and mRNA sequencing with Novogene (Cambridge, UK). FASTQ files were aligned to the human reference genome GRCh38 (Ensembl release 113) and transcript quantification performed using Salmon. All analyses including differential gene expression and functional enrichment were performed in R (version 4.5.1)^35^ (**Supplemental Figure 3, See Supplemental Methods**). All genes mapped to the Y chromosome were excluded from analysis due to the KOLF2 material transfer agreement.

### Protein studies

Batch-stored iPSC-CMs protein pellets were processed for Western blotting using the NuPAGE™ 3 to 8% Tris-Acetate Mini Protein Gel (Invitrogen™, EA03755BOX) and nitrocellulose dry membrane transfer using the iBlot™ system (Invitrogen™, IB301001) and probed with anti-*SVIL* or β-tubulin control (**See Supplemental Methods**). Cell pellets were sent for bulk proteomics analysis by the Berlin Institute of Health (BIH) Proteomics core unit. Downstream analyses including differential protein expression and functional enrichment analyses were performed in R (version 4.5.1)^35^ (**Supplemental Figure 4, See Supplemental Methods**).

### Statistical analyses

Statistical analyses were performed using GraphPad Prism (version 11) and R (version 4.5.1). Where indicated after normality testing, parametric testing was conducted using one-way analysis of variance (ANOVA) followed by Dunnett or Tukey multiple comparisons test. Non-parametric analyses were performed using the Kruskal–Wallis test with Dunn’s post hoc test for multiple comparisons. *P*<0.05 was determined to be statistically significant. For RNA-seq and proteomic analyses, Benjamini-Hochberg correction was applied for multiple comparisons, with a false discovery rate (FDR)- adjusted p-value (padj) <0.05 and absolute log2 fold change >1 defined as statistically significant differentially expressed genes (DEGs) and proteins (DEPs).

## Results

### *SVIL* LoF variants cause haploinsufficiency of supervillin in iPSC derived cardiomyocytes

A patient specific LoF variant Glu255Ter (Q255X) located in exon 6 of *SVIL* was identified as a HCM risk variant from a previous rare variant study^2^ (**Figure 1A**). This LoF variant affects the major isoforms of *SVIL* including *SVIL*-201 (**Figure 1B**) and shows expression in the heart and muscle in GTEx^9^ (**Supplemental Figure 1**). Supervillin is a known actin interacting protein and immunofluorescence (IF) confirmed colocalization of supervillin with the sarcomeric Z-disk and nucleus with no visual localization across the sarcomeric actin band outside of the discrete area of the Z-disk (**Figure 1C and D**). Having established the expression and localization of supervillin in iPSC-CMs we used CRIPSR/Cas-9 with a HDR strategy to create the isogenic heterozygous *SVIL*^Q255X/+^ and homozygous *SVIL*^Q255X/Q255X^ variant lines mimicking LoF variants observed in patients (**Figure 1E**). Screening for chromosomal abnormalities and pluripotency confirmed the stability of these cell lines (**Supplemental Figure 5**). Prediction of off-target edits was performed and the top 5 predicted off-targets were sequenced showing no off-target editing (**Supplemental Table 2 & Supplemental Figure 6**). These cell lines were differentiated into iPSC-CMs (**Supplemental Figure 7**). RT-qPCR analysis of iPSC-CMs from *SVIL*^Q255X/+^ (mean±SEM: 0.477±0.067, p=5.6×10^−6^)) and *SVIL*^Q255X/Q255X^ (0.259±0.038, p=6×10^−10^) variant lines showed a sequential allele specific reduction in RNA abundance compared to WT (0.921±0.062) (**Figure 1F**), which translated into protein level haploinsufficiency assessed by western blotting (WT=1.1±0.13; *SVIL*^Q255X/+^=0.3±0.1, p=4×10^−5^; *SVIL*^Q255X/Q255X^ =0.008±0.002, p=1×10^−6^) (**Figure 1G and Supplemental Figure 8**). IF images taken on WT, *SVIL*^Q255X/+^ and *SVIL*^Q255X/Q255X^ variant iPSC-CMs showed a qualitative dose-dependent reduction in supervillin localization in the sarcomere of the iPSC-CMs (**Figure 1H**). Additionally bulk RNA sequencing of iPSC-CMs showed a dose-dependent reduction in *SVIL* RNA (predominantly *SVIL*-201 but also seen across other transcripts) with differential transcript usage (DTU) analysis confirming *SVIL*-201 as the predominant transcript in iPSC-CMs with no evidence of isoform switching occurring alongside LoF variant inheritance (**Supplemental Figure 3C and 9**).

**Figure 1:**
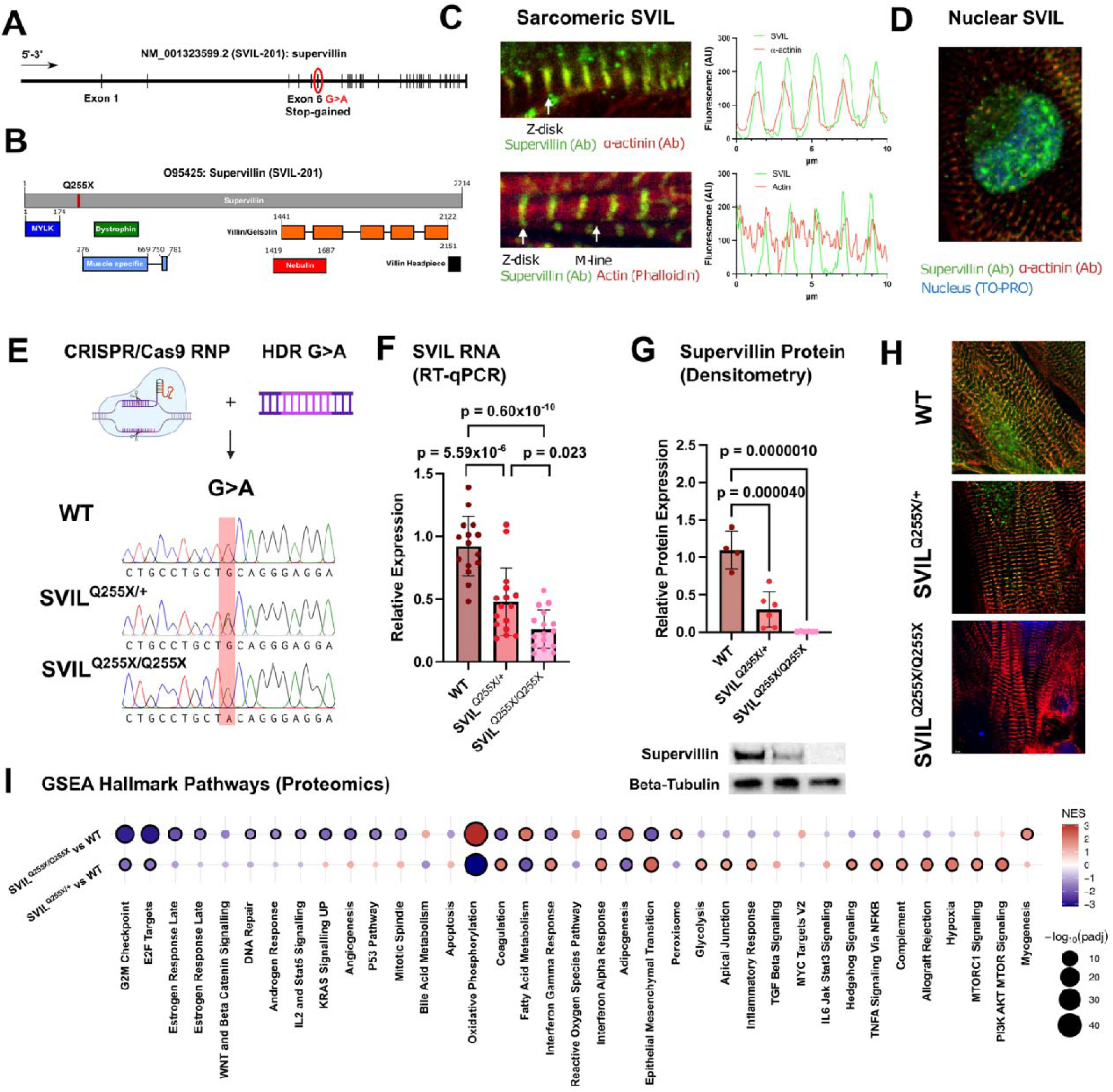
*SVIL* LoF variants cause haploinsufficiency in iPSC-CMs with divergent Hallmark pathways across heterozygous and homozygous LoF. **(A)** Schematic of the *SVIL*-201 transcript showing the Q255X variant site. **(B)** Supervillin protein showing the interaction domains with other proteins and the location of the Q255X variant. **(C)** Immunofluorescence images of iPSC-CMs stained for Supervillin, Alpha-actinin, and Actin with fluorescence shown across sarcomeres showing the localization of each protein. **(D)** iPSC-CMs stained for nuclei using TO-PRO and Supervillin and alpha-actinin b antibody showing localization of Supervillin independent of alpha-actinin at the nucleus. **(E)** Schemati showing the successful CRISPR/Cas-9 engineering of the Q255X variant as a heterozygous and homozygous G>A substitutions by Sanger sequencing. **(F)** RT-qPCR for *SVIL* shows RNA nonsense mediated decay across *SVIL*^Q255X/+^ and *SVIL*^Q255X/Q255X^ iPSC-CMs (n = 16 biological replicates). **(G)** Western blots of supervillin protein show allelic dose-dependent reduction across *SVIL*^Q255X/+^ and *SVIL*^Q255X/Q255X^ iPSC-CMs (n = 6 biological replicates). **(H)** Immunofluorescence showing qualitative sequential loss of supervillin by antibody (green) within iPSC-CMs with *SVIL*^Q255X/+^ and *SVIL*^Q255X/Q255X^ mutations. Alpha-actinin antibody shown in red and nuclear stain by TO-PRO (blue). **(I)** Gene set enrichment analysis (GSEA) of LC/MS proteomics of WT, *SVIL*^Q255X/+^, and *SVIL*^Q255X/Q255X^ iPSC-CMs (n = 6 biological replicates per genotype) showing Hallmark pathways with those reaching statistical significance to WT circled bold (FDR-adjusted p-value <0.05).

### Supervillin LoF variants drive disturbances in cellular electrophysiology, calcium handling and myosin head conformations

To understand the impact of *SVIL* LoF on iPSC-CM cellular function WT, *SVIL*^Q255X/+^, and *SVIL*^Q255X/Q255X^ lines were differentiated and stained with FluoVolt, transduced with RGECO-AV, or alpha-actinin-GFP-AV to measure cellular action potentials, calcium transients, and contractility (**Figure 2A**). iPSC-CMs were stained with FluoVolt showing localized cell membrane signal (**Figure 2B**) which displayed characteristic action potential waveforms (**Figure 2C**). The normalised and averaged action potential waveforms were acquired for *SVIL*^Q255X/+^ and *SVIL*^Q255X/Q255X^ iPSC-CMs (**Figure 2D**). Action potential (AP) duration was unaffected between genotype groups: WT (mean±SEM: 354.5±5.40 ms), *SVIL*^Q255X/+^ (336.6±7.64 ms, p=0.11) and *SVIL*^Q255X/Q255X^ line (340.3±6.65 ms, p=0.22) (**Figure 2E**). Time for transient to reach peak (AP T_on_) was significantly reduced across both *SVIL*^Q255X/+^ (28.64±0.62 ms, p=3.1×10^−4^) and *SVIL*^Q255X/Q255X^ (28.03±0.35 ms, p=9.3×10^−7^) variant lines when compared to WT (32.55±0.86 ms) (**Figure 2F**). Time of 50% transient decay (AP T_50off_) was shorter in *SVIL*^Q255X/+^ (139.5±4.32 ms, p=3.3×10^−5^), and *SVIL*^Q255X/Q255X^ (102.4±3.57 ms, p=1.2×10^−15^) when compared to WT (163.2±3.50 ms) iPSC-CMs (**Figure 2G**). Calcium transient analysis was performed using RGECO-AV transduction of iPSC-CMs in combination with automated CalTrack analysis (**Figure 2H-M**). iPSC-CMs were able to be transduced robustly and uniformly with the RGECO-AV construct (**Figure 2H**). Fluorescent imaging allowed the capture of reproducible calcium transient waveforms under 1.5 Hz pacing (**Figure 2I**). Normalized and averaged calcium transients show waveform shapes across WT, *SVIL*^Q255X/+^, and *SVIL*^Q255X/Q255X^ variant lines (**Figure 2J**). Calcium duration accounts for the length of the entire calcium transient which was longer in *SVIL*^Q255X/+^ (503.9±1.678 ms, p=0.017) compared to WT (496.6±1.92 ms) and unaffected in *SVIL*^Q255X/Q255X^ iPSC-CMs (494.7±2.29 ms, p=0.73) (**Figure 2K**). The time for the calcium transient to reach its peak (Calcium T_on_) was longer in *SVIL*^Q255X/+^ vs WT (113.9±1.22 ms vs 107.7±1.05 ms, p=9.1×10^−5^), and unaffected in *SVIL*^Q255X/Q255X^ (106.9±0.97 ms, p=0.83) (**Figure 2L**). The fit for the calcium transient decay (Tau) was longer in both *SVIL*^Q255X/+^ (283.1±5.09 ms, p=1.6×10^−15^), and *SVIL*^Q255X/Q255X^ (231.1±4.61 ms, p=0.0092) variant lines compared to WT (212.5±4.23 ms) (**Figure 2M**).

**Figure 2:**
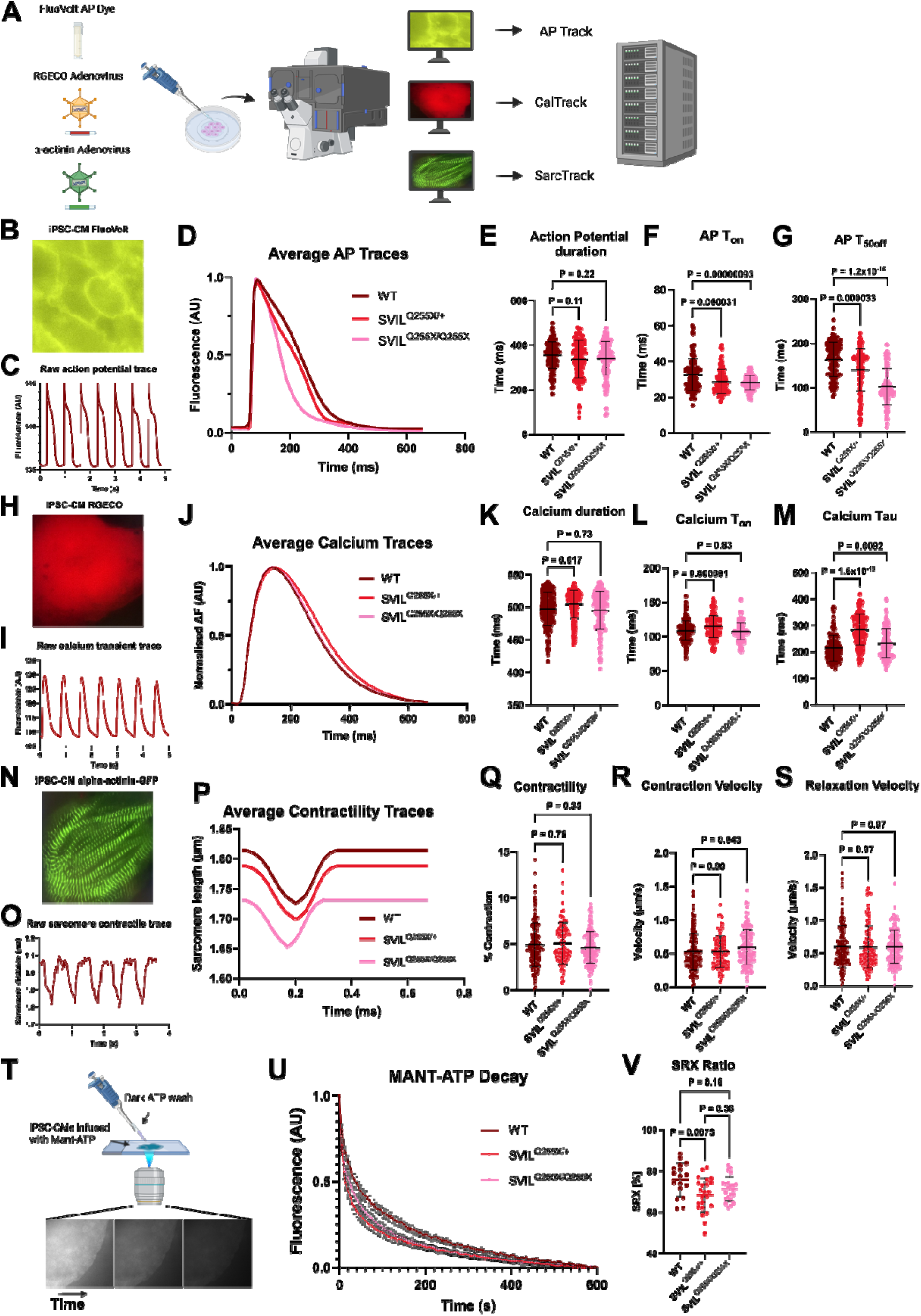
*SVIL* LoF variants alter action potential and calcium transient kinetics without affecting contractility, with heterozygous LoF altering myosin SRX/DRX ratios. **(A)** Schematic showing the three indicators and analysis pipelines used for action potentials, calcium transients, and contractility in live cell fluorescent imaging. **(B)** Representative image from movie stack of iPSC-CMs stained using FluoVolt action potential dye. **(C)** Extracted action potential (AP) traces from an AP imaging stack covering sequential iPSC-CMs depolarisations under electrical pacing (1.5 Hz). **(D)** Normalized and averaged AP traces for the entire dataset across WT (n = 123), *SVIL*^Q255X/+^ (n = 124), and *SVIL*^Q255X/Q255X^ iPSC-CMs (n = 129 technical replicates across 4 biological replicates per genotype). **(E)** Dot plot of AP duration across WT, *SVIL*^Q255X/+^, and *SVIL*^Q255X/Q255X^ iPSC-CMs. Bars showing mean ± standard deviation. **(F)** Dot plot of AP time to reach peak (T_on_) across WT, *SVIL*^Q255X/+^, and *SVIL*^Q255X/Q255X^ iPSC-CMs. Bars showing mean ± standard deviation. **(G)** Dot plot of AP time to reach 50% decay (T_50off_) across WT, *SVIL*^Q255X/+^, and *SVIL*^Q255X/Q255X^ iPSC-CMs. Bars showing mean ± standard deviation. **(H)** Representative image from movie stack of iPSC-CMs transduced with RGECO calcium transient indicator. **(I)** Extracted calcium transient traces from a calcium transient image stack covering sequential calcium transients elicited by electrical pacing of iPSC-CMs (1.5 Hz). **(J)** Normalized and averaged calcium transients for the entire dataset across WT (n = 188), *SVIL*^Q255X/+^ (n = 171), and *SVIL*^Q255X/Q255X^ iPSC-CMs (n = 169 total technical replicates, 4 biological replicates per genotype). **(K)** Dot plot of the calcium transient duration across WT, *SVIL*^Q255X/+^, and *SVIL*^Q255X/Q255X^ iPSC-CMs. Bars showing mean ± standard deviation. **(L)** Dot plot of the time to reach calcium transient peak (Calcium T_on_) across WT, *SVIL*^Q255X/+^, and *SVIL*^Q255X/Q255X^ iPSC-CMs. Bars showing mean ± standard deviation. **(M)** Dot plot of the calcium transient decay constant (tau) across WT, *SVIL*^Q255X/+^, and *SVIL*^Q255X/Q255X^ iPSC-CMs. Bars showing mean ± standard deviation. **(N)** Representative image from movie stack of iPSC-CMs stained using transduction of alpha-actinin GFP to visualise the z-disks of sarcomeres. **(O)** Extracted sarcomere contractile traces from a contractility image stack covering sequential beats elicited by electrical pacing of iPSC-CMs (1.5 Hz). **(P)** Normalized and averaged simulated contractility traces for the entire dataset across WT (n = 156 technical replicates across 5 biological replicates), *SVIL*^Q255X/+^ (92 technical replicates across 4 biological replicates), and *SVIL*^Q255X/Q255X^ iPSC-CMs (181 technical replicates across 5 biological replicates). **(Q)** Dot plot of sarcomere level contractility across WT, *SVIL*^Q255X/+^, and *SVIL*^Q255X/Q255X^ iPSC-CMs. Bars showing mean ± standard deviation. **(R)** Dot plot of contraction velocity across WT, *SVIL*^Q255X/+^, and *SVIL*^Q255X/Q255X^ iPSC- CMs. Bars showing mean ± standard deviation. **(S)** Dot plot of relaxation velocity across WT, *SVIL*^Q255X/+^, and *SVIL*^Q255X/Q255X^ iPSC-CMs. Bars showing mean ± standard deviation. Outliers were removed using the Robust regression and Outlier removal (ROUT) method with Q = 1%. (**T**)Schematic showing the experimental set-up for the Mant-ATP chase experiment with three images of a representative decay of Mant fluorescence during dark-ATP wash. **(U)** Averaged and normalized Mant-ATP decay curves across WT (n = 17), *SVIL*^Q255X/+^ (n = 23), and *SVIL*^Q255X/Q255X^ iPSC-CMs (n = 22). Each point indicates a mean ± standard error of the mean, where data is fit by a double exponential decay. **(V)** Calculated myosin SRX as a percentage of total myosin across WT, *SVIL*^Q255X/+^, and *SVIL*^Q255X/Q255X^ iPSC-CMs. Bars showing mean ± standard deviation.

Next, 2D contractility was assessed using the alpha-actinin GFP-AV in combination with SarcTrack (**Figure 2N**). Individual sarcomere tracking showed a robust contractile waveform that could be automatedly detected using the SarcTrack algorithmic identification of alpha-actinin GFP labelled sarcomeres (**Figure 2O**). Average contractility traces show the sarcomere length of contracting sarcomeres across WT, *SVIL*^Q255X/+^, and *SVIL*^Q255X/Q255X^ lines (**Figure 2P**). Fractional sarcomere shortening was measured from peak systole and peak diastole which remained unchanged between genotype groups: *SVIL*^Q255X/+^ (5.06±0.23 %, p=0.76) and *SVIL*^Q255X/Q255X^ (4.6±0.13 %, p=0.35) compared to WT (4.89±0.19%). (**Figure 2Q**). Contraction velocity calculated from the peak sarcomere shortening as a function of time showed an increased contraction velocity in *SVIL*^Q255X/Q255X^ iPSC-CMs (0.59±0.02 µm/s, p=0.04) when compared to WT (0.52±0.02 µm/s) (**Figure 2R**). Relaxation velocity measured as the percentage of sarcomeric relaxation per unit time remained unchanged between genotype groups (**Figure 2S**). To confirm this result in a more physiologically relevant loaded tissue system we used EHTs formed from WT, *SVIL*^Q255X/+^, and *SVIL*^Q255X/Q255X^ iPSC-CMs which had no measurable changes in contractile function (**Supplemental Figure 11**).

The Mant-ATP chase assay (**Figure 2T**) was used to understand if *SVIL* LoF variants impacted myosin biochemical states (**Figure 2U-V**). Averaged fluorescence decay curves show that *SVIL*^Q255X/+^ exhibits on average a faster decay when compared to WT and *SVIL*^Q255X/Q255X^ (**Figure 2U**). This translates into a significantly reduced assumed proportion of super relaxed myosin (SRX) in *SVIL*^Q255X/+^ vs WT (68.3±1.7 % vs 75.8±1.9 %, p=0.0073), which is not altered in *SVIL*^Q255X/Q255X^ iPSC-CMs (71.3±1.3 %, p=0.16) (**Figure 2V**).

### *SVIL*^Q255X/+^ causes sarcomere level hypertrophy, disorganization, and cellular stress triggering fibrotic signalling pathways alongside altered nuclear morphology

To understand if *SVIL* LoF alters contractile machinery structure, we employed alpha-actinin GFP AV to label sarcomeres and myofibrils (**Figure 3A**) in combination with SarcAsM analysis for assessing contractile morphology (**Figure 3B- K**). A custom version of SarcAsM used a TIFF frame of a relaxed iPSC-CM from each movie stack (**Figure 3B**) to extract sarcomere domains (**Figure 3C**), myofibril lines (**Figure 3D**), and myofibril lengths (**Figure 3E**). The mean transverse length of each Z- disk was measured showing increased lengths in *SVIL*^Q255X/+^ (2.283±0.06 µm, p=1.2×10^−7^) and *SVIL*^Q255X/Q255X^ (2.225±0.03 µm, p=8.1×10^−8^) when compared to WT (1.91±0.04 µm), inferring wider sarcomeres (**Figure 3F**). Acute treatment using 1 µM mavacamten for 1 hour (**Figure 3L**) showed rescue of transverse Z-disk length in *SVIL*^Q255X/+^ (Δ distance 1.02±0.03 µm, p=0.94) and partial rescue in *SVIL*^Q255X/Q255X^ (Δ distance 1.08±0.02 µm, p=0.016) when compared to WT (**Figure 3M)**. Z-disk operational order parameter (OOP) was used to measure sarcomere order where 1 defines fully aligned sarcomeres and 0 defines randomly ordered sarcomeres, *SVIL*^Q255X/Q255X^ (0.506±0.016, p=0.0051) showed a reduced OOP when compared to WT (0.577±0.016) (**Figure 3G**). Z-disk straightness measurements showed that both *SVIL*^Q255X/+^ (0.793±0.003, p=4.4×10^−8^) and *SVIL*^Q255X/Q255X^ (0.794±0.002, p=2.0×10^−9^) had altered Z-disk alignment – a hallmark of sarcomeric disarray – with mavacamten rescue in *SVIL*^Q255X/+^ (Δ1.001±0.004, p=0.99) and partial rescue in *SVIL*^Q255X/Q255X^ (Δ0.9908±0.002, p=0.002) (**Figure 3N**). Relaxed sarcomere length (SL) was measured for all groups and showed a reduced SL for *SVIL*^Q255X/+^ (1.787±0.009 µm, p=0.0094) and *SVIL*^Q255X/Q255X^ (1.731±0.005 µm, p=2.3×10^−13^) compared to WT (1.814±0.005 µm), with mavacamten increasing SL for both *SVIL*^Q255X/+^ (1.992±0.158 µm, p=8×10^−15^) and *SVIL*^Q255X/Q255X^ (1.862±0.004 µm, p=2.5×10^−8^) (**Figure 3I and Supplemental Figure 12**). Myofibril length was found to be longer in both *SVIL*^Q255X/+^ (21.60±0.61 µm, p=3.4×10^−4^) and *SVIL*^Q255X/Q255X^ (21.85±0.36 µm, p=2.0×10^−6^) compared to WT (19.09±0.37 µm) providing a signal of sarcomeric hypertrophy (**Figure 3J**). Mavacamten treatment rescued myofibril length in both *SVIL*^Q255X/+^ (Δ1.004±0.02, p=0.999) and *SVIL*^Q255X/Q255X^ (Δ0.957±0.02, p=0.39) (**Figure 3O**). We next assessed the average number of sarcomeres per myofibril finding that both *SVIL*^Q255X/+^ (12.08±0.33, p=5.7×10^−5^) and *SVIL*^Q255X/Q255X^ (12.64±0.21, p=4.0×10^−11^) had a greater number of sarcomeres per myofibril compared to WT (10.54±0.20 (**Figure 3K**). Taken together these results insinuate a degree of myofibril disorganisation and hypertrophy across both *SVIL*^Q255X/+^ and *SVIL*^Q255X/Q255X^. We further investigated nuclear morphology of *SVIL* iPSC-CMs given supervillin’s role as cytoskeletal actin interactor and found smaller nuclei area in *SVIL*^Q255X/+^ (82.1±1.33 µm^2^, p=1×10^−14^) and *SVIL*^Q255X/Q255X^ (84.24±0.93 µm^2^, p=1×10^−14^) compared to WT (112.1±1.6 µm^2^) iPSC-CMs (**Supplemental Figure 13**). Circularity as defined by 4 x (area/perimeter^2^) was also decreased in *SVIL*^Q255X/+^ (0.818±0.004, p=1×10^−14^) and *SVIL*^Q255X/Q255X^ (0.815±0.003, p=1×10^−14^) when compared to WT iPSC- CM nuclei (0.873±0.002) (**Supplemental Figure 13**).

**Figure 3:**
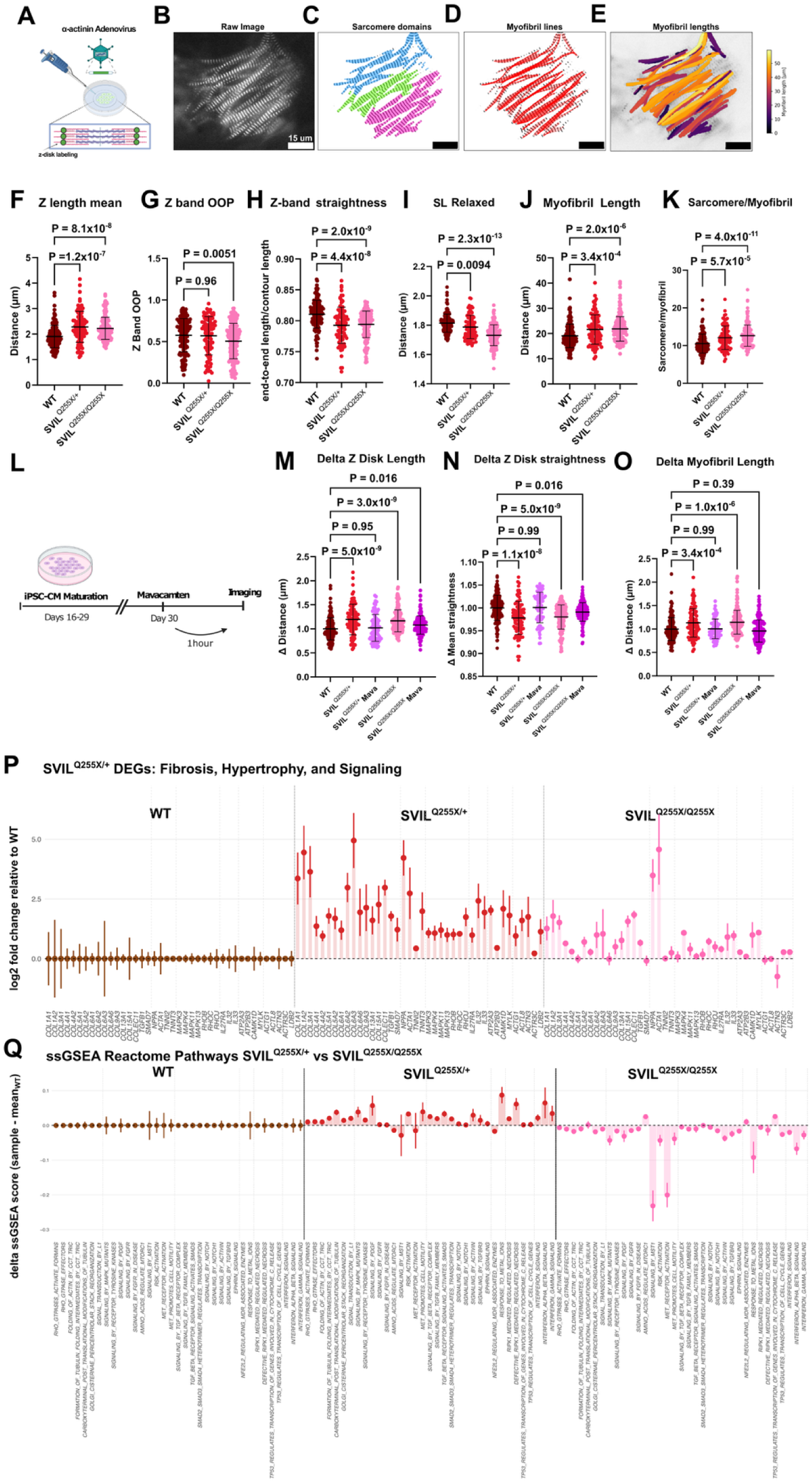
SVIL LoF variants show sarcomere level hypertrophy and disorganization, stress signalling and fibrosis marker genes. **(A)** Schematic showing the labelling of sarcomeric z-disks with alpha-actinin-GFP delivered by adenovirus. **(B)** Raw image of a relaxed iPSC-CM with alpha-actinin-GFP labelled z-disks captured by a custom SarcAsM script. **(C)** Sarcomere domains extracted from the raw image of labelled z-disks, with different colours showing nominally different sarcomere domains identified. **(D)** Myofibril lines marked by red lines showing the predicted strings of sarcomeres across an iPSC-CM. **(E)** Myofibril lengths coloured according to length of each individual string of sarcomeres identified in the myofibril lines. **(F)** Dot plot of individual transverse z-disk length calculated as the mean from each cell sampled across WT (n = 156 technical replicates across 5 biological replicates), *SVIL*^Q255X/+^ (n = 92 technical replicates across 4 biological replicates), and *SVIL*^Q255X/Q255X^ iPSC-CMs (n = 178 technical replicates across 5 biological replicates). Bars showing mean ± standard deviation for F-O. One-way ANOVA performed with post-hoc Dunnett’s multiple comparisons test. N numbers same for F-O. All parameters are in relaxed (diastolic) state. **(G)** Dot plot of z band operational order parameter (OOP) showing alignment of z-disks averaged in each iPSC-CM across WT, *SVIL*^Q255X/+^, and *SVIL*^Q255X/Q255X^ iPSC-CMs. **(H)** Dot plot of sarcomere level z-disk straightness averaged per iPSC-CM across WT, *SVIL*^Q255X/+^, and *SVIL*^Q255X/Q255X^ iPSC-CMs. **(I)** Dot plot of relaxed average sarcomere length (SL) per iPSC-CM across WT, *SVIL*^Q255X/+^, and *SVIL*^Q255X/Q255X^ iPSC-CMs. **(J)** Dot plot of myofibril length averaged per iPSC-CM across WT, *SVIL*^Q255X/+^, and *SVIL*^Q255X/Q255X^ iPSC-CMs. **(K)** Dot plot showing the average number of sarcomeres per myofibril per iPSC-CM across WT, *SVIL*^Q255X/+^, and *SVIL*^Q255X/Q255X^ iPSC-CMs. **(L)** iPSC-CMs were incubated with 1uM mavacamten or DMSO (vehicle) control for 1 hour prior to imaging**. (M**) Dot plot of mavacamten treatment showing delta mean transverse Z disk length per iPSC- CM with *SVIL*^Q255X/+^ + mavacamten (n = 76) and *SVIL*^Q255X/Q255X^ + mavacamten (n = 150) compared to WT iPSC-CMs. N numbers same M-O. **(N)** Dot plot of mavacamten treatment showing delta mean z-disk straightness per iPSC-CM per genotype group compared to WT. **(O)** Dot plot of mavacamten treatment showing delta mean myofibril length per iPSC-CM per genotype group compared to WT. **(P)** Lollipop plot of RNA significant DEGs between the *SVIL*^Q255X/+^ vs WT contrast (*SVIL*^Q255X/Q255X^ for comparison) under themes of fibrosis/ECM remodelling, hypertrophic/foetal gene reprogramming, MAPK/Rho signalling, interleukin/inflammatory signalling, and cytoskeletal genes. Bars show mean ± SEM of sample-level Log2 fold change relative to WT of DESeq2 size-factor normalized counts. **(Q)** Lollipop plot of proteomic single sample GSEA (ssGSEA) scores for Reactome pathways showing statistically significant differences between *SVIL*^Q255X/+^ and *SVIL*^Q255X/Q255X^. Pathways are sub-grouped into themes of cytoskeleton and mechanotransduction, hypertrophic signalling, fibrosis, oxidative stress, cell death pathways, and interferon signalling. Bars show mean ± SEM of the delta ssGSEA scores (sample – mean_WT_). N = 6 biological replicates per genotype.

Alongside sarcomere level effects of *SVIL* LoF we found that *SVIL*^Q255X/+^ variants showed a variety of DEGs consistent with RNA signatures of pro-fibrosis, hypertrophy, inflammatory signalling, and structural remodelling (**Figure 3P**). Except for *NPPA* and *ACTA1,* these DEGs were not conserved in *SVIL*^Q255X/Q255X^ variant iPSC-CMs (**Figure 3P**). Analysis focusing on long non-coding RNAs (lncRNAs) found four shared and upregulated DEGs between the mutant cell lines, of which lncRNA-H19 is implicated in cardiac fibrosis and hypertrophy^36^ (**Supplemental Figure 10**). Furthermore, delta ssGSEA analysis of Reactome pathway related proteins showed that *SVIL*^Q255X/+^ and *SVIL*^Q255X/Q255X^ iPSC-CMs showed divergent signatures of signalling involving fibrosis and mechanotransduction, hypertrophy, ROS pathways, cell death, and interferon based inflammatory signalling (**Figure 3Q**). Taken together, this showed RNA and protein level evidence for HCM signatures in *SVIL*^Q255X/+^ that are not present in *SVIL*^Q255X/Q255X^. This was further reinforced by over-representation analysis (ORA) of GO-BP terms associated with HCM specifically, where there was an enrichment of disease specific terms only in *SVIL*^Q255X/+^ (**Supplemental Figure 14**). We also assessed concordance between all RNA and protein level changes **(Supplemental Figure 15).**

### *SVIL* heterozygous and homozygous LoF variants exhibit divergent energy metabolism traits

We used gene set enrichment analysis (GSEA) of Hallmark pathways performed on LC/MS proteomic data of *SVIL*^Q255X/+^ and *SVIL*^Q255X/Q255X^ iPSC- CMs as an unbiased top-down approach to investigate energy metabolism. This showed contrasting Hallmark pathway responses (namely oxidative phosphorylation (OXPHOS), fatty acid metabolism, interferon alpha and gamma response, epithelial mesenchymal transition (EMT), adipogenesis and coagulation pathways) across heterozygous and homozygous variant iPSC-CMs (**Figure 1I**). Hallmark GSEA analysis in *SVIL*^Q255X/+^ identified significant positive enrichment of hypoxia (NES=2.01, FDR=4.76×10^−7^), glycolysis (NES=1.48, FDR=0.0089), TNFα signalling via NFKB (NES=1.79, FDR=2.54×10^−4^), interferon alpha (NES=1.99, FDR=5.18×10^−4^), interferon gamma (NES=1.89, FDR=2.03×10^−4^), complement (NES=1.68, FDR=0.0023), EMT (NES=2.32, FDR=3.74×10^−9^), coagulation (NES=1.92, FDR=3.26×10^−4^), MTORC1 (NES=1.68, FDR=4.23×10^−4^) and PI3K/AKT/MTOR (NES=2.12, FDR=2.01×10^−5^) pathways. By contrast, these alterations were not significant or were negatively enriched in *SVIL*^Q255X/Q255X^ iPSC-CMs (**Figure 1I**). Notably, OXPHOS showed the strongest negative enrichment in *SVIL*^Q255X/+^ (NES= -3.17, FDR=1.3×10^−36^) contrasted by a strong positive enrichment in *SVIL*^Q255X/Q255X^ iPSC-CMs (NES=3.09, FDR=1.32×10^−41^).

To further investigate the mechanism behind the divergent metabolic phenotype seen in GSEA Hallmark Pathways between the mutant cell lines, we stained iPSC-CMs to visualise mitochondria. This qualitatively showed a decrease in mitochondria in *SVIL*^Q255X/+^ iPSC-CMs compared to WT and *SVIL*^Q255X/Q255X^ iPSC-CMs (**Figure 4A-C**). We further used Seahorse flux analysis to assay iPSC-CM OCR (**Figure 4D**). Normalized OCR is shown for WT, *SVIL*^Q255X/+^, and *SVIL*^Q255X/Q255X^ iPSC-CMs using the Seahorse assay (**Figure 4E**). Basal respiration was decreased in *SVIL*^Q255X/+^ iPSC-CMs (25.33±3.74 pmol/min) when compared to WT (51.27±5.77 pmol/min, p=0.048) and *SVIL*^Q255X/Q255X^ iPSC-CMs (65.6±3.03 pmol/min, p=2×10^−4^) and unaffected between *SVIL*^Q255X/Q255X^ and WT iPSC-CMs (p=0.36) (**Figure 4F**). Maximal respiration was decreased in *SVIL*^Q255X/+^ (66.37±12.07 pmol/min) when compared to WT (221.5±21.43 pmol/min, p=0.004) and *SVIL*^Q255X/Q255X^ iPSC-CMs (173.8±11.96 pmol/min, p=0.02) and unaffected between *SVIL*^Q255X/Q255X^ and WT iPSC-CMs (p=0.73) (**Figure 4G**). ATP- linked respiration decreased in *SVIL*^Q255X/+^ (22.0±3.41 pmol/min) when compared to *SVIL*^Q255X/Q255X^ iPSC-CMs (58.97±2.7 pmol/min, p=1×10^−4^) but there was no change between *SVIL*^Q255X/+^ and WT iPSC-CMs (39.0±4.1 pmol/min, p=0.17) (**Figure 4H**). Proton leak was decreased in *SVIL*^Q255X/+^ (3.33±0.44 pmol/min) when compared to WT (12.27±2.49 pmol/min, p=0.0002) and *SVIL*^Q255X/Q255X^ iPSC-CMs (6.63±0.5 pmol/min, p=0.036) (**Figure 4I**). Spare capacity was decreased in *SVIL*^Q255X/+^ (41.04±8.69 pmol/min) when compared to WT iPSC-CMs (170.3±17.07 pmol/min, p=1×10^−4^) which was not significant when compared to *SVIL*^Q255X/Q255X^ iPSC-CMs (108.2±10.95 pmol/min, p=0.065) (**Figure 4J**). Mavacamten treatment for 24 hours at 1 µM did not rescue OCR and other flux analysis parameters in *SVIL*^Q255X/+^ (**Supplemental Figure 16**). RNAseq DEGs showed a consistent profile of energetic dysregulation and GO-BP showed hypoxia and ECM related pathways in *SVIL*^Q255X/+^ that were not evident in *SVIL*^Q255X/Q255X^ iPSC-CMs (**Figure 5K Supplemental Figure 17**). Particularly, the RNAs involved in the hypoxic inducible factor (HIF) mediated hypoxia related metabolic switch were upregulated in *SVIL*^Q255X/+^ (**Figure 4K**), consistent with a profile of pseudohypoxia in *SVIL*^Q255X/+^ iPSC-CMs. Moreover, delta ssGSEA analysis using Reactome pathway related proteins showed divergence in mitochondrial metabolism (including tricarboxylic acid cycle (TCA) and OXPHOS) and fatty acid metabolism pathways between *SVIL*^Q255X/+^ and *SVIL*^Q255X/Q255X^ iPSC-CMs (**Figure 4L**).

**Figure 4:**
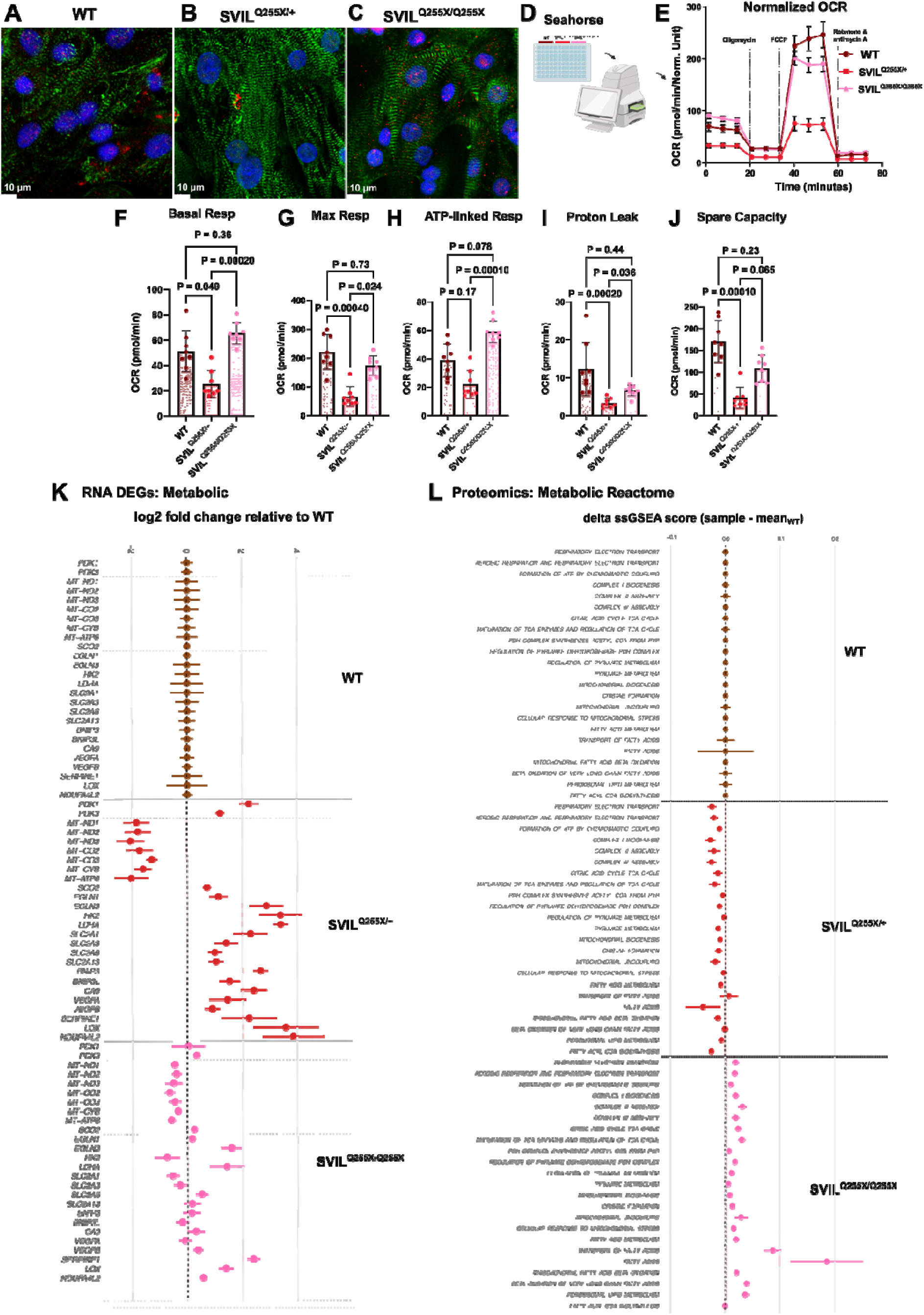
SVIL heterozygous LoF variants alter OCR and mitochondrial metabolic pathways showing a signature of pseudohypoxia. (**A-C**) Representative immunofluorescence images of (**A**) WT, (**B**) SVIL^Q255X/+^, and (**C**) SVIL^Q255X/Q255X^ iPSC-CMs stained with mitochondrial anti-ATP5A antibody. **(D)** Schematic showing the plate format used to set-up for Seahorse flux analysis of WT, *SVIL*^Q255X/+^, and *SVIL*^Q255X/Q255X^ iPSC-CMs. **(E)** Averaged and normalized oxygen consumption rates measured across WT, SVIL^Q255X/+^, and SVIL^Q255X/Q255X^ iPSC-CMs. N = 8 technical replicates for each genotype group (for plots E-J) with trends confirmed over 2 biological replicates. Each point is plot as mean ± standard deviation. **(F)** Dot plot of basal respiration per well sampled across WT, SVIL^Q255X/+^, and SVIL^Q255X/Q255X^ iPSC-CMs. Bars showing mean ± standard deviation. **(G)** Dot plot of maximal respiration per well sampled across WT, SVIL^Q255X/+^, and SVIL^Q255X/Q255X^ iPSC-CMs. Bars showing mean ± standard deviation. **(H)** Dot plot of ATP-linked respiration per well sampled across WT, SVIL^Q255X/+^, and SVIL^Q255X/Q255X^ iPSC-CMs. Bars showing mean ± standard deviation. **(I)** Dot plot of proton leak per well sampled across WT, SVIL^Q255X/+^, and SVIL^Q255X/Q255X^ iPSC-CMs. Bars showing mean ± standard deviation. **(J)** Dot plot of spare capacity per well sampled across WT, SVIL^Q255X/+^, and SVIL^Q255X/Q255X^ iPSC-CMs. Bars showing mean ± standard deviation. **(K)** Lollipop plot of metabolism linked RNA significant DEGs between the *SVIL*^Q255X/+^ vs WT contrast (*SVIL*^Q255X/Q255X^ vs WT for comparison) grouped by pyruvate dehydrogenase kinases (PDKs), oxidative phosphorylation, and HIF-mediated glycolytic switch. Bars show mean ± SEM of sample-level log2 fold change relative to WT of DESeq2 size-factor normalized counts. **(L)** Lollipop plot of proteomic single sample GSEA (ssGSEA) scores for Reactome pathways showing statistically significant differences between *SVIL*^Q255X/+^ and *SVIL*^Q255X/Q255X^. Pathways are sub-grouped into themes of mitochondrial function/metabolism, and fatty acid/lipid metabolism. Bars show mean ± SEM of the delta ssGSEA scores (sample – mean_WT_). N = 6 biological replicates per genotype.

**Figure 5:**
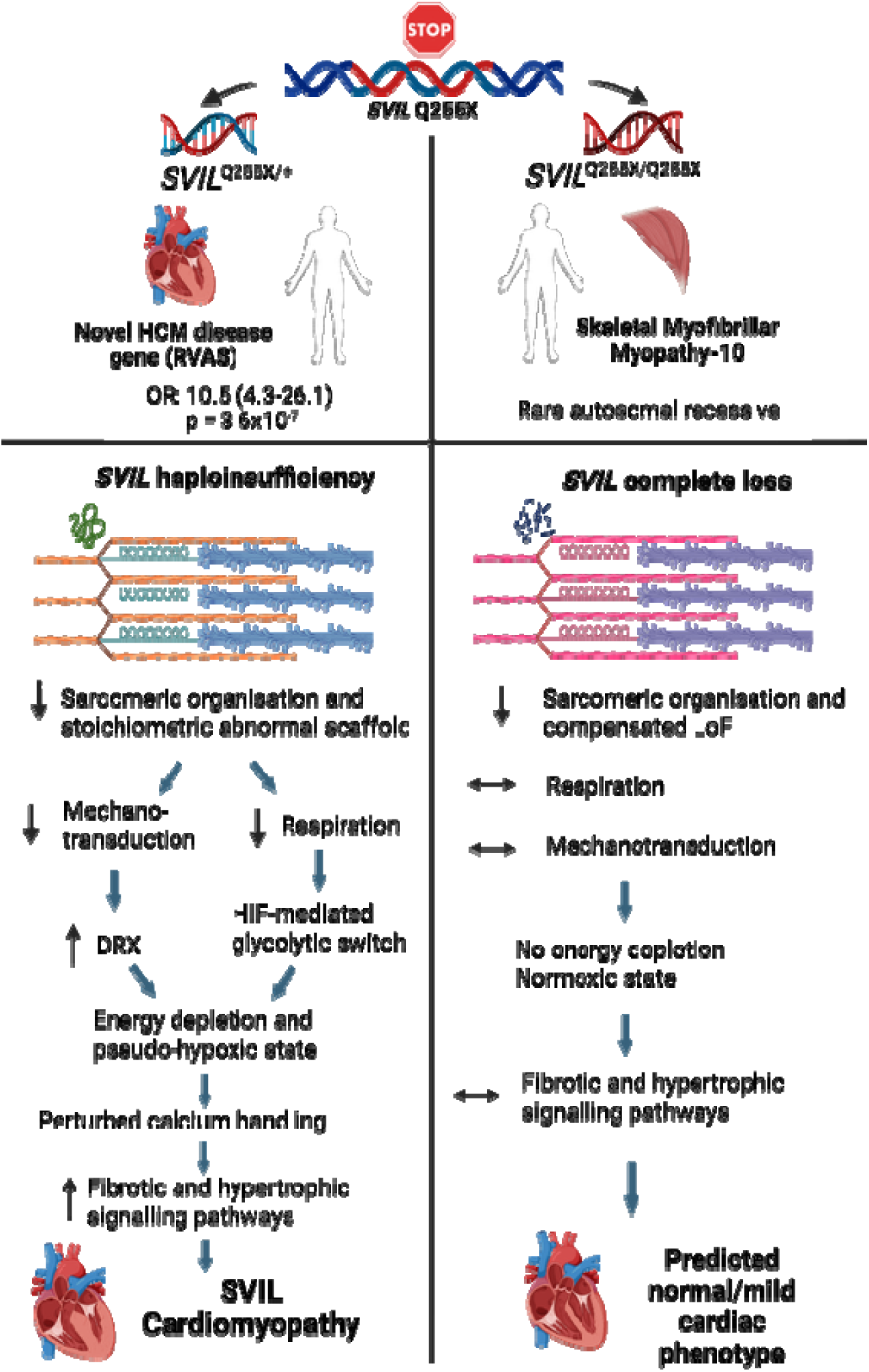
SVIL Paradigm Overview: The SVIL-201 transcript encodes supervillin in the adult human heart. A rare variant study has discovered SVIL heterozygous loss of function variants as a novel driver of HCM. Homozygous SVIL loss of function variants have been found to exhibit a skeletal myofibrillar myopathy-10, with little evidence of a strong cardiac phenotype.

## Discussion

Using a patient specific human iPSC-CM disease model of supervillin cardiomyopathy, we demonstrated for the first time a novel disease mechanism driven by variants in a non-canonical actin-interacting Z-disk and costameric scaffolding gene (**Figure 5**). The key findings are: *SVIL* truncating variants cause HCM via haploinsufficiency; *SVIL* LoF heterozygosity is more deleterious than homozygosity in cardiomyocytes; sarcomeric disorganization and mitochondrial dysfunction drive disease via impaired mechanotransduction and cellular respiration, respectively; pseudohypoxia is a key disease mechanism that drives metabolic remodelling; and mavacamten rescues sarcomeric disorganization but not mitochondrial dysfunction.

We established that the patient specific Q255X variant in *SVIL* caused an allelic dose-dependent RNA decrease in iPSC-CMs consistent with nonsense mediated decay and protein level haploinsufficiency as validated by RT-qPCR (**Figure 1F**), western blotting (**Figure 1G**), and corroborated by multi-omics analysis (**Supplemental Figure 3C and 4C**). Importantly, we did not find evidence of a truncated supervillin protein in our LC/MS analysis. This is unique in that most variants that drive HCM are missense and dominant negative except for those in *MYBPC3*^24^ and *ALPK3*^4^, and missense variants in Z-disk genes such as *FLNC* commonly cause HCM whereas truncating LoF variants cause dilated cardiomyopathy (DCM)^6^. For this reason, gene therapies that focus on *SVIL* cardiomyopathy upregulation are attractive.

Supervillin haploinsufficiency is further unique in that heterozygous variant carrying iPSC-CMs displayed features consistent with HCM whereas homozygous variant iPSC-CMs did not. This is consistent with the limited existing clinical literature^2,20^ and more insight into phenotypes of homozygous carriers would be illuminating; however, this is limited by their rarity. Four homozygous *SVIL* LoF patients reported in Hedberg-Oldfors et. al.^20^ displayed with the novel skeletal MFM-10 with mild cardiac features consistent with Z-disk related myopathies^6^. Interestingly, current (unpublished) clinical evidence does not suggest features of skeletal myopathy in heterozygous variant carriers. One could postulate that the predominant skeletal myopathy in homozygous carriers could be because supervillin is a direct nebulin interactor in skeletal muscles, where nebulin variants are a major driver of Nemaline myopathy^37^. Existing literature describes supervillin as a key actin interactor that is important in myofibrillogenesis, the nuclear lamina, and the costamere, and that the loss of this protein within the myocyte may interfere with contractile function or ability to translocate force^8,10,11^. Interestingly, our mutant iPSC-CMs showed an allelic dose-dependent decrease in sarcomere length suggesting a hypercontracted basal state with supervillin loss (**Figure 2P and 3I**). This could be explained by supervillin acting as an additional linkage between the actin filaments and the sarcolemma at costameres as proposed by Oh et. al.^8^, of which LoF would result in impaired tethering of the sarcomeres to the sarcolemma and thus impaired intra- and intercellular mechanotransduction.

Further, partial and complete loss of this important scaffolding protein is evidenced by the increased sarcomeric disorganisation and smaller nuclei in mutant iPSC-CMs which directly implicate an impaired process of mechanotransduction. This bidirectional mechanotransduction occurs predominantly at costameres and focal adhesions (and internally via Z-disks) which also act as mechanosensors that activate downward stress and hypertrophic signalling pathways^38^. We hypothesize that in *SVIL* LoF, partial loss as seen in *SVIL*^Q255X/+^ is more disruptive than complete loss, as abnormal and mixed complexes at these mechanotransducers alter stoichiometric balance which is more deleterious than complete absence. It is possible that heterozygous *SVIL* truncation preserves enough protein to misassemble the force-transmission complexes altering signalling thresholds and triggers a downstream stress and hypertrophic signalling response, whereas homozygous *SVIL* LoF behaves as a compensated LoF state. Our multi-omics findings add weight to the dysregulated mechanotransduction hypothesis in *SVIL*^Q255X/+^ as we observe increased fibrosis, hypertrophic signalling pathways (MTORC1, PI3K/AKT/MTOR and MAPK/Rho), and inflammatory response in *SVIL*^Q255X/+^ that are absent in *SVIL*^Q255X/Q255X^ iPSC-CMs (**Figures 1I**, **3P and 3Q**). This finding was mirrored when we interrogated HCM linked GO-BP terms across *SVIL*^Q255X/+^ and *SVIL*^Q255X/Q255X^ iPSC-CMs, where *SVIL*^Q255X/+^ showed a variety of key HCM pathways as enriched that were not present in *SVIL*^Q255X/Q255X^ iPSC-CMs (**Supplemental Figure 14**). Interestingly, the lncRNA-H19 is upregulated in pathological cardiac hypertrophy^36^ and we found evidence of this in both *SVIL*^Q255X/+^ and *SVIL*^Q255X/Q255X^ iPSC-CMs, with the homozygous state displaying a greater log2-fold change (**Supplemental Figure 10**). LncRNA-H19 is a negative regulator of cardiac hypertrophy via the calcium/calmodulin-dependent protein kinase IIδ and NFAT pathways and has complex roles in fibrosis, angiogenesis and inflammatory response^36^. Taken together, this suggests a degree of compensation in both mutant cell lines and lends support to the compensated homozygous LoF state hypothesis.

In contrast to variants in core sarcomeric genes which cause HCM through inefficient sarcomeric ATP utilization resulting in increased energy demand and leading to cellular energy depletion^22^, *SVIL*^Q255X/+^ directly perturbs mitochondrial function showing reduced OCR and ETC respiratory capacity that was not evident in *SVIL*^Q255X/Q255X^ iPSC-CMs as validated by our IF staining, Seahorse, and multi-omics findings. It is possible that supervillin functions like the intermediate filament desmin, another mechanotransducing Z-disk and costameric protein that produces desminopathies (another type of myofibrillar myopathy with cardiac features) in deficient states. Desmin deficiency is well known for causing mitochondrial dysfunction in striated muscle via various mechanisms such as mitochondrial mislocalization^39^, and desmin KO mice hearts showed decreased mitochondrial numbers, ultrastructural defects and severely impaired OXPHOS and fatty acid metabolism^40^. Since desmin is not a predicted supervillin interactor on StringDB nor a DEG or DEP in our omics data and together with upregulated mitophagy genes *BNIP3* and *BNIP3L* in *SVIL*^Q255X/+^ suggesting increased mitochondrial turnover, we hypothesize that there could be a similar mechanism for mitochondrial dysfunction in *SVIL* cardiomyopathy.

We further substantiated the energetic findings by exploring the DEGs associated with metabolic pathways and found increased expression of pyruvate dehydrogenase kinases (PDKs) and genes related to glucose metabolism and transport, as well as decreased expression of genes involved in the mitochondrial electron transport chain (ETC) in *SVIL*^Q255X/+^ iPSC-CMs (**Figure 4K**). PDK1 phosphorylates and inhibits pyruvate dehydrogenase (PDH) which prevents pyruvate conversion into acetyl-CoA, and lactate dehydrogenase A (LDHA) catalyses conversion of pyruvate to lactate, which together divert pyruvate away from the TCA cycle and towards lactate production^41^. Along with other increased canonical HIF-signalling targets such as VEGFA, SERPINE1, and NDUFA4L2, the likely mechanism is a HIF-mediated glycolytic switch due to a pseudohypoxic cellular state in *SVIL*^Q255X/+^ iPSC-CMs that is not present in *SVIL*^Q255X/Q255X^ iPSC-CMs. This is corroborated with transcriptomic ORA using GO-BP which displays a strong theme of cellular response to hypoxia in *SVIL*^Q255X/+^ iPSC-CMs (**Supplemental Figure 17**). Proteomic delta ssGSEA analysis of Reactome pathways further substantiates this showing negatively enriched pathways relating to fatty acid metabolism, pyruvate metabolism, PDH complex, TCA cycle, mitochondrial complexes, aerobic respiration and electron transport in *SVIL*^Q255X/+^ iPSC-CMs that are opposite and enriched in *SVIL*^Q255X/Q255X^ iPSC-CMs (**Figure 4L**).

Moreover, we believe that the decreased SRX in *SVIL*^Q255X/+^ is secondary and not a primary disease driver. Unlike classical hypercontractile sarcomeric HCM variants that destabilize the myosin mesa and the IHM^42,43^, the reduced SRX and increased DRX proportions in *SVIL*^Q255X/+^ were not linked to increased contractility or increased cellular energetics. We hypothesize that myosin may be affected as a secondary compensatory effect to maintain contractility that occurs alongside the impaired mechanotransduction in the supervillin haploinsufficiency state. The secondary increase in DRX in *SVIL*^Q255X/+^ further drives increased energy demand in an already energy depleted cardiomyocyte. Thus, this pseudohypoxic cellular state results in supply/demand mismatch in subcellular compartments such as the SERCA2 pump^22^ resulting in prolonged calcium transients as seen in *SVIL*^Q255X/+^ iPSC-CMs (**Figure 2J-M**), which translates clinically to diastolic dysfunction. This leads to hypertrophic signalling by the calcium-calmodulin-dependent protein kinase (CaMK) and MAPK pathways evidenced by our multi-omics findings.

Lastly, we found that the allosteric myosin inhibitor mavacamten rescued sarcomeric disarray and reduced fractional shortening in both mutant lines (**Figure 3M-O**, **Supplemental Figure 12**) but 24-hour treatment did not rescue the mitochondrial dysfunction in *SVIL*^Q255X/+^ iPSC-CMs as it has done previously in classical sarcomeric HCM^23^ (**Supplemental Figure 16**). Mavacamten is known to reduce hypercontractility in thick filament HCM by increasing the SRX ratio^23,24^ and rescues calcium hypersensitivity and dysregulation in thin filament HCM^28^. However, given that *SVIL* cardiomyopathy belongs to the group of Z-disk/costameric HCM, mavacamten usage in this group is not directly targeted at the pathomechanism of disease. Since the increased DRX in *SVIL*^Q255X/+^ iPSC-CMs is potentially compensatory, reversal of this could create an acute systolic insufficiency. However, it will be important to know if mitochondrial dysfunction in *SVIL* haploinsufficiency can be rescued using chronic mavacamten treatment through its positive effects on sarcomeric organization. Therapies targeting mitochondrial bioenergetics may represent a novel therapeutic strategy for *SVIL*-associated cardiomyopathy alongside other existing modalities.

## Conclusions

We demonstrate in a human iPSC-CM patient model the mechanisms that drive HCM in *SVIL* LoF variants validating previously published rare variant association findings. We propose a novel pathomechanism of haploinsufficiency in a scaffolding protein that alters mechanotransduction and cellular energetics triggering hypertrophic and pro-fibrotic signalling cascades driving a HCM phenotype. We establish a mechanistic framework of *SVIL*-associated HCM confirming its importance as a HCM gene.

## Acknowledgments

This work was supported by a Wellcome Sir Henry Dale Fellowship (222567/Z/21/Z) and a BHF CRE Award (RE/24/130024) to C.N.T. The Oppenheimer Memorial Trust Graduate Scholarship and the Keith Murray Graduate Scholarship of Lincoln College supported Y.J.L. This work was also supported by the British Heart Foundation’s Big Beat Challenge award to CureHeart H.W. (award reference no. BBC/F/21/220106) and Oxford CRE3 (RE/18/3/34214). The BHF Research Grant Award (FS/4yPhD/F/21/34157) supported IB. The RDM Scholars Programme supported S.TM.J. N.B. is participant in the BIH Charité Clinician Scientist Program funded by the Charité – Universitätsmedizin Berlin and the Berlin Institute of Health at Charité. J.S.W. was supported by Medical Research Council (UK), Sir Jules Thorn Charitable Trust [21JTA], British Heart Foundation [RE/24/130023; BBC/F/21/220106], and the NIHR Imperial Biomedical Research Centre. BR was supported by Oxford NIHR Biomedical Research Centre and funded by a Wellcome Career Development Award fellowship (302210/Z/23/Z). BR also acknowledges support from the Oxford British Heart Foundation Centre of Research Excellence.

The views expressed in this work are those of the authors and not necessarily those of the funders.¬¬

For the purpose of open access, the authors have applied a Creative Commons Attribution (CC BY) licence to any Author Accepted Manuscript version arising.

## Non-standard Abbreviations and Acronyms

HCM: Hypertrophic Cardiomyopathy
HCMSARC+: Sarcomeric HCM
HCMSARC-: Non-Sarcomeric HCM
LoF: Loss of Function
GWAS: Genome Wide Association Study
MFM: Myofibrillar Myopathy
GTEx: Genotype-Tissue Expression
HDR: Homology-Directed Repair
DEG: Differentially Expressed Gene
DEP: Differentially Expressed Protein
LC/MS: Liquid Chromatography – Mass Spectrometry
AV: Adenovirus
GSEA: Gene Set Enrichment Analysis
ssGSEA: Single sample GSEA
ORA: Over-representation Analysis
GO-BP: Gene Ontology – Biological Process
IHM: Interacting Heads Motif
SRX: Super Relaxed State
DRX: Disordered Relaxed State
OXPHOS: Oxidative Phosphorylation
MAPK: Mitogen-Activated Protein Kinase
SERCA: Sarcoplasmic/endoplasmic reticulum calcium-ATPase

