## Supplemental_FIle for "Unmasking Supervillin*: SVIL* haploinsufficiency causes hypertrophic cardiomyopathy by impairing mechanotransduction and cellular energetics"

**Short title**: *SVIL variants drive a novel mechanism for HCM*

**Authors:** Yifan J. Li[1], MBChB, BMedSci (Hons); Yiangos Psaras[1,2, †], DPhil, MSc, BSc (Hons); Violetta Steeples[1, †], DPhil, BSc (Hons); Josephine M. Watkins[1,3], BSc; Charlotte Hooper[1], PhD, BSc; Marta Moya-Jódar[1,2], PhD, MSc, BSc; Thomas Nicol[4], PhD, BbMedSci; Alexander J. Sparrow[1], PhD, MSc, BSc (Hons); Marcos Garcia-Lacarte[5], PhD, MSc, BSc; Samuel TM Jones[1], MBiol; Isabelle Bond[6], MSc, BSc (Hons); Niklas Beyhoff[1,7], MD; Paul Robinson, DPhil, BSc (Hons); Marieluise Kirchner[8], PhD; Philipp Mertins[8], PhD; James S Ware[10,11,12], PhD, FMedSci; R Thomas Lumbers[9], MB Bchir, PhD; Betty Raman[1], MBBS, DPhil; Hugh Watkins[1,5], FRS, FMedSci; Christopher N. Toepfer[1,2*], PhD, BSc (Hons).

[†]Joint Second Author

*Corresponding author

**This PDF includes:** **Supplemental Methods**

**Supplemental Tables 1-5**

**Supplemental Figures 1-17**

**Supplemental References**

### Supplemental Methods

#### Generation of SVIL Q255X iPSC lines using CRISPR/Cas-9

The KOLF2 line was reprogrammed by Human iPSC Initiative (HipSci) as previously described[1]. Custom single-stranded guide-RNA (crRNA) and HDR template were obtained from Integrated DNA Technologies (IDT) (**Supplemental Table 3**). Nucleofection was performed using the Lonza P3 Primary Cell 4D Nucleofector Kit (Lonza, V4XP-3024) on the Lonza Nucleofector (Lonza, AAF-1002B) using 6uL gRNA (5 µL of crRNA annealed with 5 µL TracrRNA) and 3uL Alt-R s.p. HiFi Cas9 nuclease (forming the ribonucleoprotein (RNP) complex) and HDR template, as previously described[2]. The Non-Homology End-Joining (NHEJ) inhibitor Alt-R™ HDR Enhancer V2 (IDT, 10007921) was used to improve efficiency. PCR primers were designed using PrimerBlast[3] and optimised. DNA extraction and PCR amplification was done using Phire Tissue-Direct PCR Kit (ThermoFisher Scientific, F170S) and Sanger sequencing performed by Source BioScience (Cambridge, UK). Two rounds of clonal selections were done to obtain isogenic wild-type (WT), heterozygous ($\mathit{SVIL}^{\mathrm{Q255X/+}}$) and homozygous ($\mathit{SVIL}^{\mathrm{Q255X/Q255X}}$) lines and ensure monoclonality. Karyotyping abnormalities were screened using the hPSC Genetic Analysis Kit (StemCell Technologies, 07550) (**Supplemental Figure 5A)**. Pluripotency was verified using immunofluorescence (IF) (StemLight™ Pluripotency Transcription Factor Antibody Kit, 9093) and quantitative real-time PCR (RT-qPCR) (TaqMan™ FAM probes, Applied Biosystems Hs02387400_g1, Hs00742896_s1, Hs00602736) (**Supplemental Figure 5B**). The top five predicted off-target edits by CRISPOR[4] were screened using PCR and Sanger sequencing (**Supplemental Table 2 and Supplemental Figure 6**).

#### iPSC culture and CM differentiation

iPSCs were cultured in E8™ Flex (ThermoFisher Scientific, A2858501) and propagated on 1% Geltrex (ThermoFisher Scientific, A1413302)-coated 6-well plates incubated at 37°C with 5% $CO_2$. Passaging was done at 80% confluency using 0.5mM EDTA in PBS into fresh E8™ Flex with 10 µM of ROCK-inhibitor Y-27632 (Selleck, #S1049) for 24 hours. CM differentiations were done using a published protocol[5] with modifications using RPMI-1640 with N21-MAX (Biotechne, AR008) supplemented with 65 µg/mL L-ascorbic acid (LAA) instead of B27[6] (**Supplemental Figure 7A**). After metabolic selection using glucose starvation, iPSC-CMs were replated between D17 and D20 using 10X TrypLE (Gibco, A12177-01) onto new plates and maintained in cardiomyocyte passaging media

(RPMI/N21-MAX +insulin + LAA, KnockOut Serum Replacement 10% (Thermofisher Scientific, 10828028), and Thiazovivin 2 µM (Selleck, S1459)) for 24 hours. Replated iPSC-CMs were used for experimentation between D30-33.

**Functional analyses of iPSC-CMs**

iPSC-CMs were replated onto 35mm Ibidi polymer coverslip dishes (Ibidi, 81156) for live-cell imaging studies using 1.8 mM calcium Tyrodes HEPES buffer **(Supplemental Table 4)** and under 100X Nikon Ti2 Eclipse microscope equipped with a Photometrics Kinetix sCMOS camera. Acute drug treatment was performed using 1 µM Mavacamten in dimethyl sulfoxide (DMSO) for 30 minutes. iPSC-CMs were paced at 1.5 Hz using bipolar pulses of 20 ms duration at 30 V. Videos were obtained at 100 fps. A single video is defined as a technical replicate and an iPSC-CM differentiation batch is defined as a biological replicate. For membrane action potential (AP), FluoVolt™ Membrane Potential Kit (ThermoFisher Scientific, F10488) was used per protocol and imaged at 252 fps after 15 minutes of incubation, and data was analysed using AP Track, a modified version of CalTrack[7]. For calcium transients and contractility, iPSC-CMs were transduced 48 hours prior imaging with replication deficient adenoviral constructs (AV) encoding RGECO (a red fluorescent calcium indicator) and α-actinin-GFP fusion protein as previously reported[7]. AP and calcium transient videos were analysed using APTrack/CalTrack[7] to quantify amplitude ($F_{Max}/F_0$), transient duration, time-to-peak ($T_{on}$), decay time ($T_{off}$), and Tau decay constant. 2D contractility videos were analysed using a modified version of SarcTrack[8] to measure contraction duration, contraction time and relaxation time. Sarcomere organisation was analysed using SarcAsM[9] to measure relaxed and contracted sarcomere length (deriving percentage contraction and velocity), Z-disk length, Z-disk orientation order parameter (OOP) (where 1 defines fully aligned sarcomeres and 0 defines randomly ordered sarcomeres), Z-disk straightness, sarcomere length, and myofibril length. This work utilized the computational resources of the NIH HPC Biowulf cluster (https://hpc.nih.gov).

**Generation and analysis of engineered heart tissues (EHTs)**

EHTs were generated using a modified protocol as published by Cumberland et. al.[5] that was adapted from the original EHT system from the Eschenhagen lab[10]. Modifications included the use of optimised media (**Supplemental Table 5**) and the use of the CuriBio Mantarray™ 24-well plates without the use of cardiac fibroblasts.

Mantarray™ lattice posts were prepared per user guide. WT, *SVIL*$^{Q255X/+}$ and *SVIL*$^{Q255X/Q255X}$ iPSC-CMs were dissociated as described and counted. Each EHT contains 1.5 million CMs. EHT master mixes were prepared on ice and 116 µL was mixed with 3.6 µL thrombin (100 U/ml) aliquot to make a single EHT by pipetting between the lattice posts of each well of the Mantarray™ 24-well plate. Casted plates were incubated at 37°C for 90 minutes before adding 500 µL of CMP into each well for a further 30-minute incubation. The lattices with EHTs attached were then carefully removed from the casting plate and transferred into newly prepared 24-well plate with EHT culture media with Thiazovivin for 24 hours, followed by normal EHT culture media for maintenance with media change on alternate days and imaged on day 21. EHTs were paced with 20V at 1 Hz and imaged under 2X Nikon Ti2 Eclipse microscope equipped with a Photometrics Kinetix sCMOS camera. Videos were analysed using BeatProfiler[11] to obtain tissue width, mean active force, mean active stress, and contraction time (**Supplemental Figure 11**).

**Seahorse mitochondrial flux analysis**

Cellular mitochondrial oxygen consumption rate (OCR) was assessed using Seahorse XF Mito Stress Test kit (Agilent, 103015-100) on the Seahorse XF Pro Analyzer per Agilent user guide. Passaged iPSC-CMs at D17 were seeded at 3000 cells/well density onto XFe96/XF Pro Cell Culture microplates and cultured until D25. For each experiment, a Seahorse XFe96 cartridge was hydrated using 200 µL per well of XF Calibrant solution (Agilent, 100840-000) one day prior experiment and kept in non-$CO_2$ incubator overnight. 0.5ml of 1M Glucose Solution (Agilent, 103577-100), 0.5ml of 100mM Pyruvate Solution (Agilent, 103578-100) and 0.5ml of 200mM Glutamine Solution (Agilent, 103579-100) were added 48.5ml of RPMI Seahorse RPMI media (Agilent, 103576-100) and warmed to 37°C. Final concentrations of compounds used were 1 µM Oligomycin, 2µM FCCP, and 0.5 µM Rotenone/Antimycin. The microplate containing iPSC-CMs were equilibrated in non-CO2 incubator for 1 hour prior experiment. Data was analysed using Seahorse Wave Desktop software and GraphPad Prism (version 11).

**RNA processing, qPCR and bulk RNA-seq**

Replated iPSC-CMs cultured to D30-33 were washed in 1X PBS, flash-frozen in liquid nitrogen and batch-stored at -80°C. RNA extraction was done using RNeasy® Mini Kit (Qiagen, 74104) with β-mercaptoethanol addition and vortex homogenisation. cDNA was generated using High-Capacity cDNA Reverse Transcription Kit (Applied Biosystems, 4368814) and qPCR performed using TaqMan™ probes for *SVIL* and β-tubulin (FAM Hs00931004_m1, VIC Hs00742828_s1) on QuantStudio 7 Flex (ThermoFisher Scientific). For RNA-Seq studies, extracted RNA samples were sent for lncRNA and mRNA sequencing with Novogene (Cambridge, UK). Sample quality control was assessed on the 5400 Fragment Analyzer System (Agilent) and all passed with RNA integrity number (RIN) of >9.7. Directional library preparation with rRNA removal was performed, and sequencing done on the NovaSeq X Plus Series (Illumina) with each sample >40M reads. FASTQ files were quality-checked and aligned to the human reference genome GRCh38 (Ensembl release 113) using the nf-core/rnasplice pipeline v1.0.4[12]. Briefly, reads were pre-processed and aligned with STAR, a splice-aware aligner. Transcript quantification was performed using Salmon, and alternative splicing analysis was conducted with the SUPPA package to estimate percent spliced-in (PSI) values across samples. All analyses were performed in R[13] (version 4.5.1) using Bioconductor[14] (version 3.21). Differential gene expression analysis was conducted using DESeq2[15] and functional enrichment analysis was performed using clusterProfiler[16] and ReactomePA[17] **(Supplemental Figure 3)**. Gene set enrichment analysis (GSEA) was performed using fgseaMultilevel[18] using the Molecular Signatures Database (MSigDB) Hallmark gene set. Differential transcript usage (DTU) was assessed by calculating within-sample transcript proportions and comparing these between genotype groups using Kruskal–Wallis tests, followed by Dunn's post hoc tests with BH correction for multiple comparisons.

**Protein extraction, Western blotting and bulk proteomics by LC-MS**

Batch-stored iPSC-CMs were lysed in radioimmunoprecipitation Assay (RIPA) buffer containing protease and phosphatase inhibitors (Roche, 04693159001, PHOSS-RO), homogenised using sonication, centrifuged (max speed, 10min at 4°C), and the supernatant used for protein quantification (Pierce™ BCA Protein Assay Kit, ThermoFisher Scientific, 23225) and prepared in NuPAGE 4X LDS Sample Buffer (ThermoFisher, NP0007) and NuPAGE™ 10X Sample Reducing Agent (ThermoFisher, NP0009). SDS-PAGE was performed using NuPAGE™ 3 to 8% Tris-Acetate Mini Protein Gels (Invitrogen™, EA03755BOX) loaded with 20 µg of lysate in each well and resolved under denaturing conditions in NuPAGE™ Tris-Acetate SDS Running Buffer (Invitrogen™, LA0041). Dry membrane transfer was performed using nitrocellulose iBlot™ Transfer Stack (Invitrogen™, IB301001) and iBlot™ 2 Gel Transfer Device (Invitrogen™, IB21001). Membranes were blocked in 5% milk/PBST (PBS + 0.1% Tween) for 40 minutes and incubated overnight in anti-*SVIL* (1:1000) or β-tubulin (Abcam, AB6046) control. After secondary antibody incubation for 1 hour between washing steps, SuperSignal™ West Dura (ThermoFisher Scientific, 34076) was applied for 5 minutes and membrane imaged using the ChemiDoc BioRad system. Batch-stored iPSC-CM pellet samples were sent for bulk proteomics analysis by the Berlin Institute of Health (BIH) Proteomics core unit. Cells were lysed in buffer containing 1% sodium deoxycholate (SDC), 150 mM NaCl, 1 mM EDTA, 10 mM DTT, 40 mM chloroacetamide, and protease inhibitor cocktail (PIC 2/3), followed by heating at 95 °C for 10 min and benzonase treatment for 20 min at room temperature. Protein (100 µg per sample) was digested overnight with trypsin at a 1:25 enzyme-to-substrate ratio, and peptides were purified using C18 solid-phase extraction columns. LC–MS/MS analysis was performed on an Orbitrap Exploris 480 mass spectrometer using a 110-min gradient in data-independent acquisition (DIA) mode with 40 variable isolation windows and 1 µg peptide injected per run. Raw data were analysed using Spectronaut (v19) against the UniProt human reference proteome (February 2024 release) with an additional contaminants database. Downstream analyses were performed in R[13] (version 4.5.1) using Bioconductor[14]. $Log_2$-transformed peptide intensities were filtered for a minimum of three valid values, MNAR-imputed, normalised, and averaged to the protein level. Differential protein abundance analysis was performed using limma[19], and enrichment analyses were performed using clusterProfiler[16] and ReactomePA[17] (**Supplemental Figure 4**). GSEA analyses on differentially expressed proteins (DEPs) were performed in similar fashion to the RNA-seq data described. Single sample GSEA (ssGSEA) analysis was performed using the Gene Set Variation Analysis (GSVA) package[20] using the MSigDB Reactome gene set, and pathway-level enrichment scores across genotypes were assessed using Kruskal-Wallis test with Dunn's post hoc tests with BH correction for multiple comparisons.

### Supplemental Tables

#### Supplemental Table 1: Antibody list

| Type | Antibody | Product code | Species | Dilution for IF |
|---|---|---|---|---|
| Primary | SVIL | Sigma (Atlas) HPA020138-25UL | Rabbit | 1:100 |
| Primary | EA-53 (alpha actinin) | Merck A7811 | Mouse | 1:100 |
| Primary | Anti-Sarcomeric Alpha Actinin | Abcam 68167 | Rabbit | 1:200 |
| Primary | ATP5A | Abcam Ab14748 | Mouse | 1:150 |
| Primary | Beta Tubulin | Abcam Ab6046 | Rabbit | N/A |
| Secondary | Alexa Fluor 488 | Invitrogen A21206 | α-rabbit | 1:200 |
| Secondary | Alexa Fluor 546 | Invitrogen A11018 | α-mouse | 1:200 |
| Secondary | To-PRO iodide 642/661 | Invitrogen T3605 | - | 1:1000 – 1:800 |
| Secondary | Rhodamine Phalloidin | Invitrogen R415 | - | 1:40 |
| Nuclear stain | NucBlue™ Live ReadyProbes™ Reagent (Hoechst 33342) | Invitrogen R37605 | - | 2 drops per 1ml |

### Supplemental Table 2: Top 5 off-target sites predicted by CRISPOR for *SVIL* Q255X

| Off target sequence | Mismatch Position | CFD score † | Chromosome | Locus Description |
|---|---|---|---|---|
| ACACAACCTCTCCCTGCAGCAGG | *....*.**........... | 0.875 | chr17 | intron:MGAT5B |
| TCACAACTCCTCTCTGCAGCTGG | *....*......*....... | 0.7 | chrX | intergenic:GS1-519E5.1-TBL1X |
| TGACAGCCCCTCCCTGCAGCAGG | **.....*............ | 0.69565217 | chr16 | intron:PRKCB |
| AGACAACTCCTCCCTGCAGCCGG | **...*.............. | 0.69565217 | chr1 | intergenic:RD3-RD3/RP11-359E8.3 |
| AAACAACTCTTCCCTGCAGCCGG | **...*...*.......... | 0.68449198 | chr11 | intergenic:FGF3-RP11-643C9.1 |

† CFD = cutting frequency determination score where 1 = high likelihood and 0 = unlikely. See Doench et. al.[21]

**Supplemental Table 3: CRISPR/Cas9 genome editing design for generation of the SVIL Q255X variant**

| Type | Sequence |
|---|---|
| gRNA | CCACAGCTCCTCCCTGCAGC |
| PAM | AGG |
| Cut to mutation distance | 0 |
| HDR template +strand | AGGGGATAGCTGTGGGTCACCAAAGGAGGGGCTCCGGGAGGCTGCCTGCTACAGGGAGGAGCTGTGGGCGTGCTTGGGGGACCGTGGCACTTCAGTGAAGG |
| HDR template -strand | CCTTCACTGAAGTGCCACGGTCCCCCAAGCACGCCCACAGCTCCTCCCTGTAGCAGGCAGCCTCCCGGAGCCCCTCCTTTGGTGACCCACAGCTATCCCCT |
| PCR forward primer | CCCTATCACACAGAGAGGCAAAA |
| PCR reverse primer | GCTGTGTGCAGTAGAGTGTCA |

**Supplemental Table 4: Modified Tyrode's HEPES solution**

| Component | Concentration (mM) |
|---|---|
| NaCl | 150 |
| HEPES | 5 |
| Glucose | 0.420 |
| KCl | 1.8 |
| NaH2PO3 | 0.350 |
| CaCl2 | 1.8 |

### Supplemental Table 5: EHT Casting Reagents

| | | |
|---|---|---|
| **Cardiomyocyte Passaging Media (CMP)** | RPMI/N21 (or B27) +insulin + LAA<br>KOSR (10%)<br>Thiazovivin (2uM)<br>Penicillin-Streptomycin | 90 ml<br>10 ml<br>20 µL<br>1ml |
| **EHT Media** | RPMI/N21 (or B27) + insulin + LAA<br>Aprotinin 33mg/ml<br>Penicillin-Streptomycin | 50 ml<br>50 µL<br>500uL |
| **Aprotinin**<br>A1153-100MG, Sigma | **33 mg/ml**<br>Aprotinin<br>Water (Filter sterilise in TC)<br>50 µl aliquots (12), labelled "A33"<br>Store -80°C | <br>100 mg<br>3.03 ml |
| **Fibrinogen**<br>F8630-1G, Sigma | **200 mg/ml**<br>Fibrinogen<br>0.9% NaCl (warmed in incubator)<br><br>Add 14.42 µl Aprotinin (33 mg/ml) | <br>1 g<br>5 ml |
| | To dissolve: warm in hands and leave on roller (can take several hours to dissolve).<br>DO NOT heat in incubator or water bath.<br><br>50 µl aliquots (20), labelled "F+A"<br>Store -80°C | |
| **Thrombin**<br>T4648-1KU, Sigma | **100 U/ml**<br>Thrombin<br>PBS (TC sterile)<br>Water (TC sterile) | <br>1000 U<br>6 ml<br>4 ml |
| | 450 µl aliquots, labelled "T100" stocks<br>**3.6 µl aliquots, labelled "T"**<br>Store -80°C | |
| **Rock Inhibitor Y-27632 (Dihydrochloride)**<br>(Abcam, Ab120129 = 1mg)<br>320.3g/mol molecular weight | **10mM** (if using 1mg, dilute 312.21 µL, if 5mg, dilute to 1.5548ml)<br>Divide into 20uL aliquots<br>Store -20°C | |
| **Geltrex™ Reduced-Growth Factor Basement-Membrane Matrix**<br>LDEV-free, stem-cell qualified (A14133-02) | **Undiluted.**<br>**60 uL** aliquots. Thaw on ice overnight. Freeze 200uL pipette tips overnight (or for 10 minutes in -80°C freezer). Pipette using frozen (pre-chilled) pipette tips into 60 uL aliquots and keep in -20°C | |
| **EHT Master Mix (1X EHT)** | **CMP (with cells)**<br>**Rock Inhibitor (10mM)**<br>**Geltrex**<br>**Fibrinogen + Aprotinin** | 100 µL<br>0.12 µL<br>12 µL<br>4 µL |

### Supplemental Figures

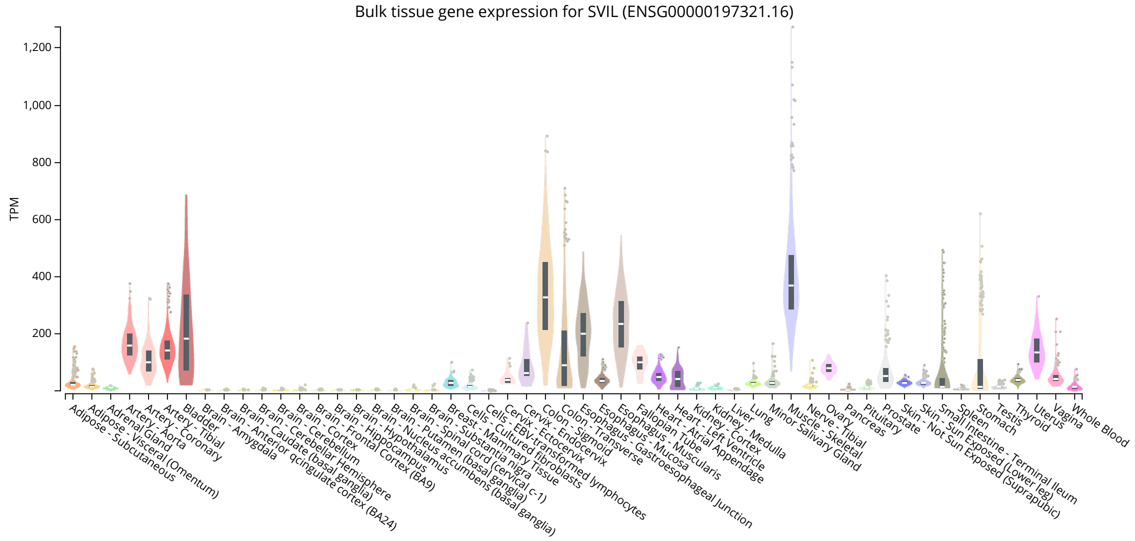


**Supplemental Figure 1: GTEx bulk tissue gene expression for *SVIL* (accessed January 2026).** Transcripts per million reported for *SVIL* across tissues of the human body per Genotype-Tissue Expression (GTEx) Portal, v10.

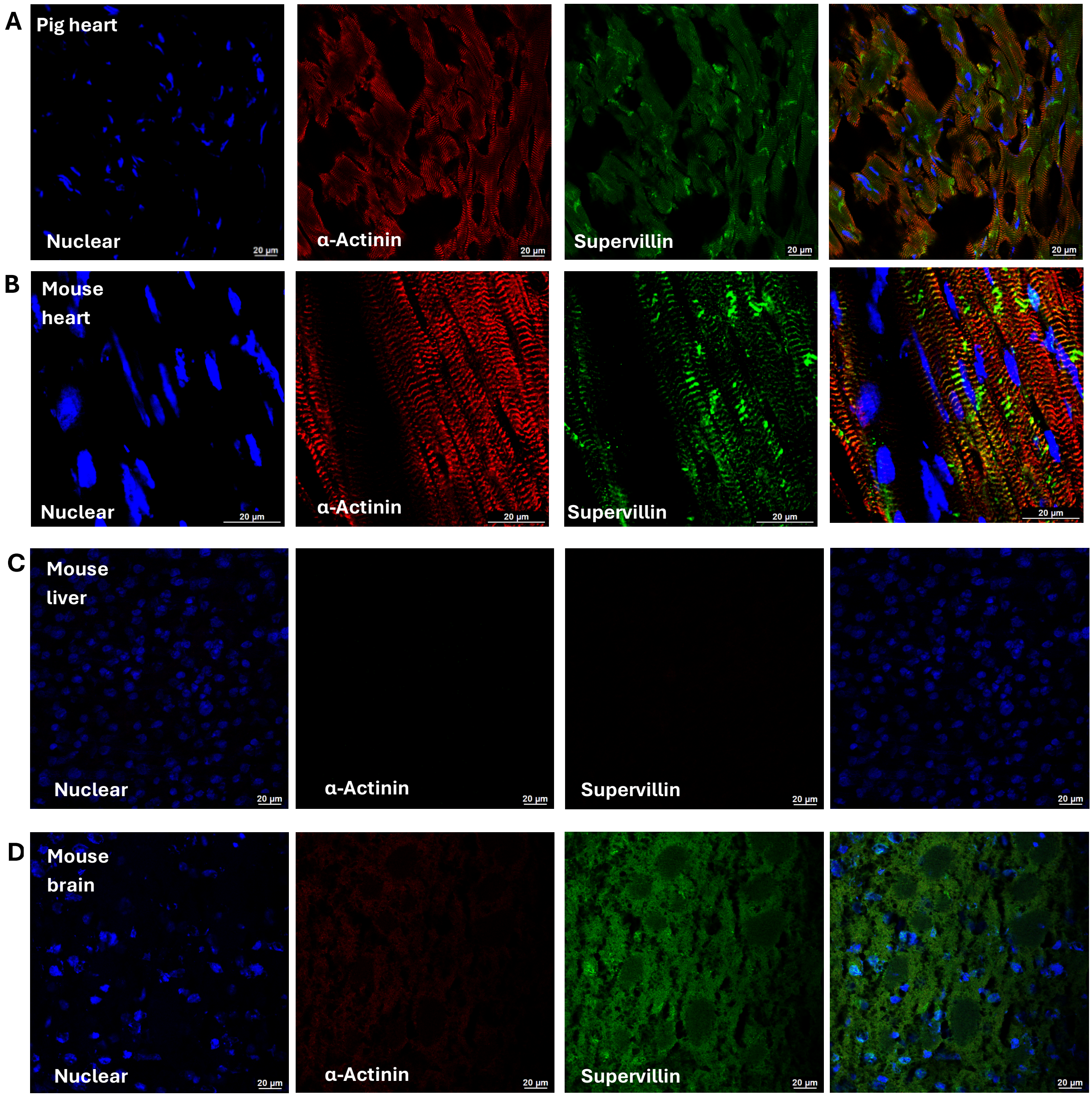


**Supplemental Figure 2: Control staining for Supervillin in pig and mouse heart show localisation at the Z-disc and mouse liver confirmed as negative control.** A) Pig left ventricular myocardium stained for nuclei using TO-PRO (Blue), alpha-actinin antibody (Red), and Supervillin antibody (Green). B) Mouse left ventricular myocardium stained for nuclei using TO-PRO (Blue), alpha-actinin antibody (Red), and Supervillin antibody (Green). C) Mouse Liver as negative control for both alpha-actinin and Supervillin stained for nuclei using TO-PRO (Blue), alpha-actinin antibody (Red), and Supervillin antibody (Green). D) Mouse brain used as negative control for alpha-actinin stained for nuclei using TO-PRO (Blue), alpha-actinin antibody (Red), and Supervillin antibody (Green). Scale bars indicate 20 μm.

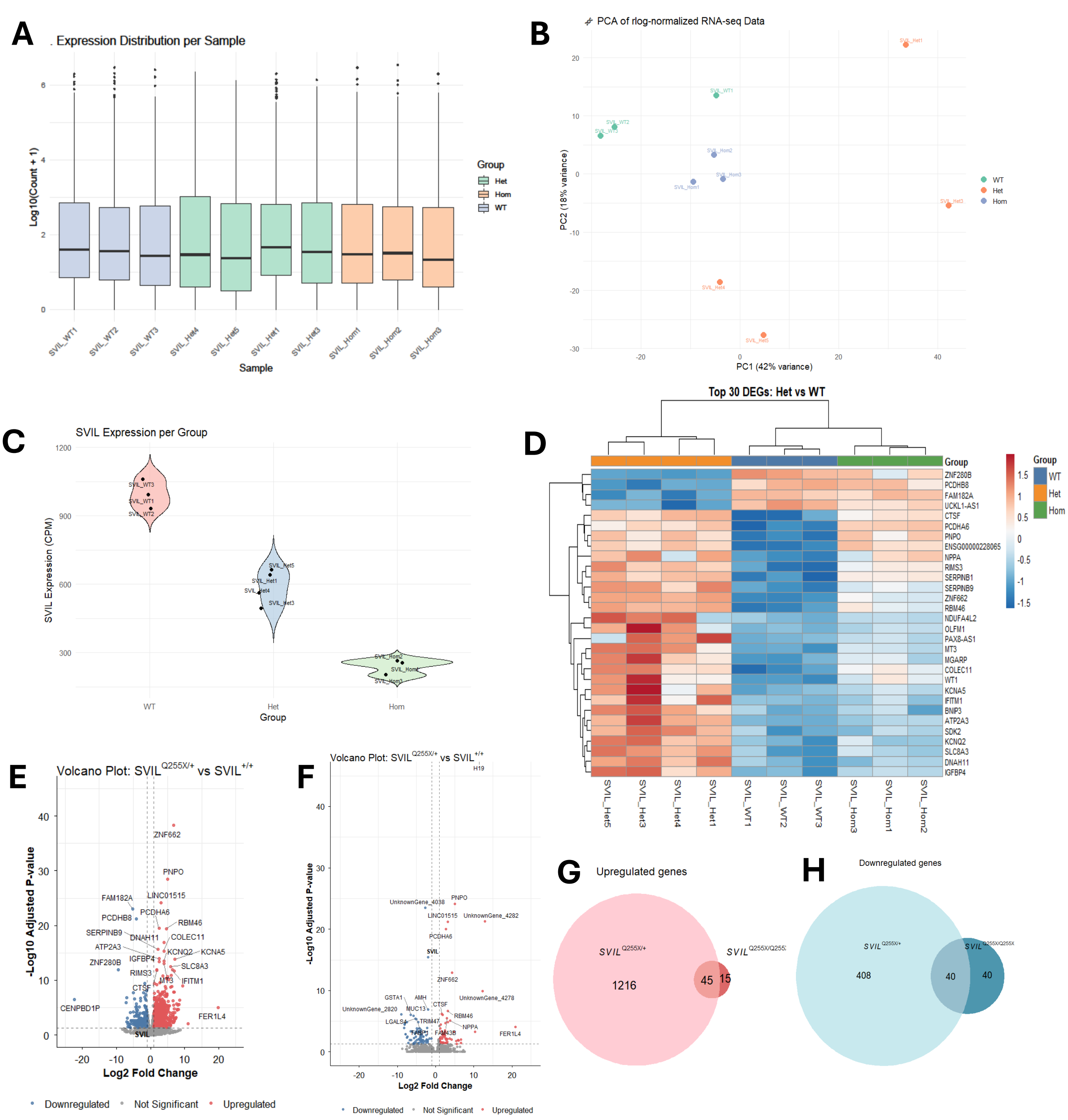


**Supplemental Figure 3: RNA-seq analysis pipeline with Differential Gene Expression (DEG) analysis. (A)** Box plot showing the log10(count+1) transformed transcriptome wide expression distribution for each sample. N = 3 for WT and SVIL$^{Q255X/Q255X}$, n = 4 for SVIL$^{Q255X/+}$. **(B)** Principal component analysis (PCA) plot showing sample clustering along PC1 and PC2. **(C)** Violin plot of SVIL expression in counts per million (CPM) per sample group. **(D)** Heatmap showing top 30 DEGs between the SVIL$^{Q255X/+}$ and WT contrast. **(E)** Volcano plot of gene expression between the SVIL$^{Q255X/+}$ and WT contrast with upregulated DEGs in red and downregulated DEGs in blue. 1709 total DEGs detected in this contrast. **(F)** Volcano plot of gene expression between the SVIL$^{Q255X/Q255X}$ and WT contrast. 140 total DEGs detected in this contrast. Log2 fold change (|log2FC|) cut off set at > 1, false discovery rate (FDR) cut off set at <0.05. **(G)** Venn diagram showing upregulated DEGs between the two contrasts with 45 overlapping genes. **(H)** Venn diagram showing downregulated DEGs between the two contrasts with 40 overlapping genes.

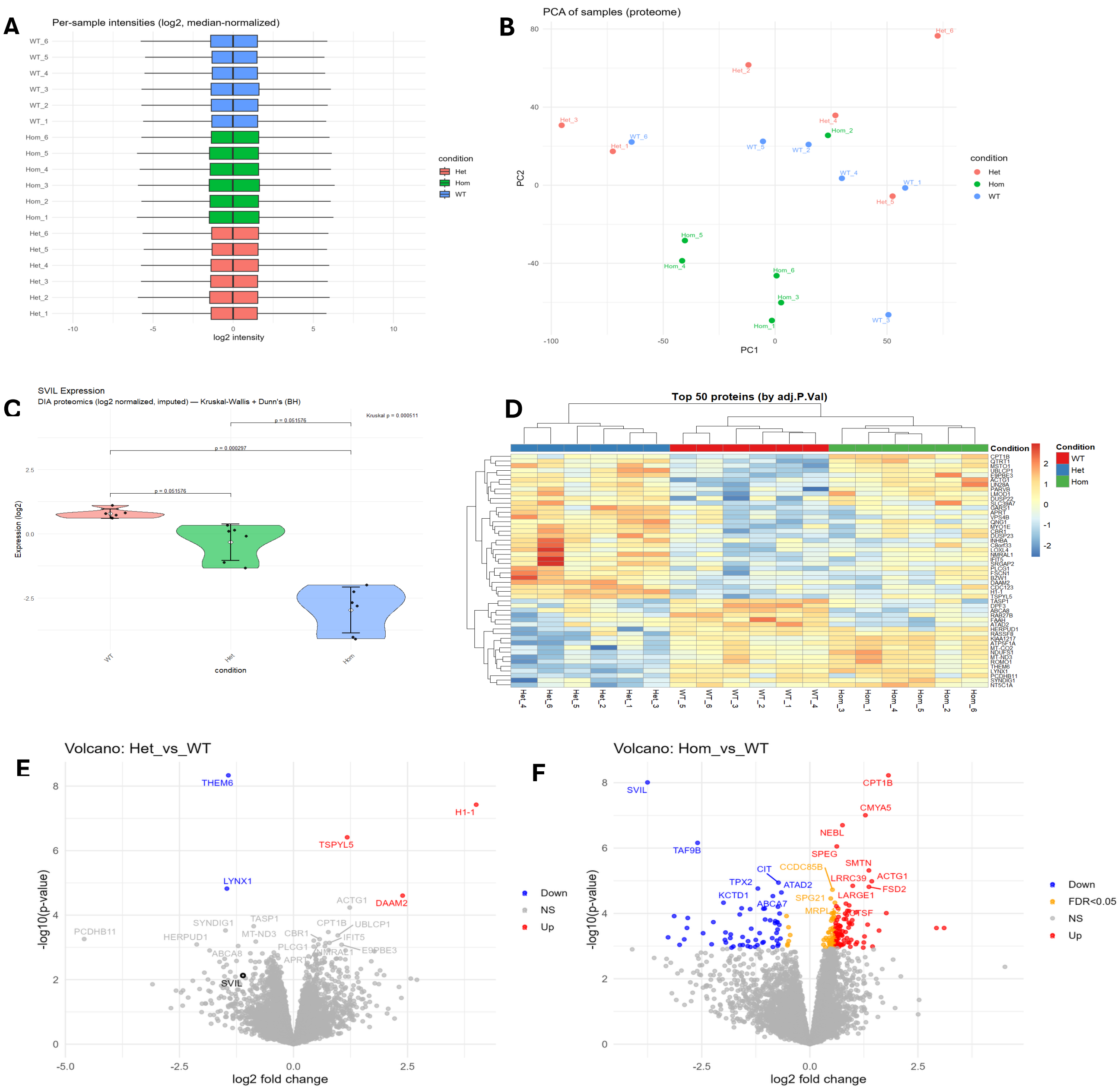


**Supplemental Figure 4: Proteomic analysis pipeline with Differential Protein Expression (DEP) analysis. (A)** Box plot of proteome wide log2 peptide intensity distribution for each sample (n = 6). **(B)** PCA plot showing sample clustering along PC1 and PC2. **(C)** Violin plot showing averaged log2 transformed SVIL peptide expression per sample group. Kruskal-Wallis test performed with post-hoc Dunn's correction. **(D)** Heatmap showing top 50 proteins ranked by FDR adjusted p-value between the SVIL$^{Q255X/+}$ and WT contrast. **(E)** Volcano plot of protein expression between the SVIL$^{Q255X/+}$ and WT contrast with upregulated DEPs in red and downregulated DEPs in blue. 5 total DEPs were detected in this contrast. **(F)** Volcano plot of protein expression between the SVIL$^{Q255X/Q255X}$ and WT contrast with upregulated DEPs in red, downregulated DEPs in blue, and proteins with FDR adjusted p-value <0.05 but log2 fold change <1 in yellow. 197 total DEPs were detected in this contrast with top 20 DEPs labelled. Log2 fold change (|log2FC|) cut off set at > 1, FDR cut off set at <0.05.

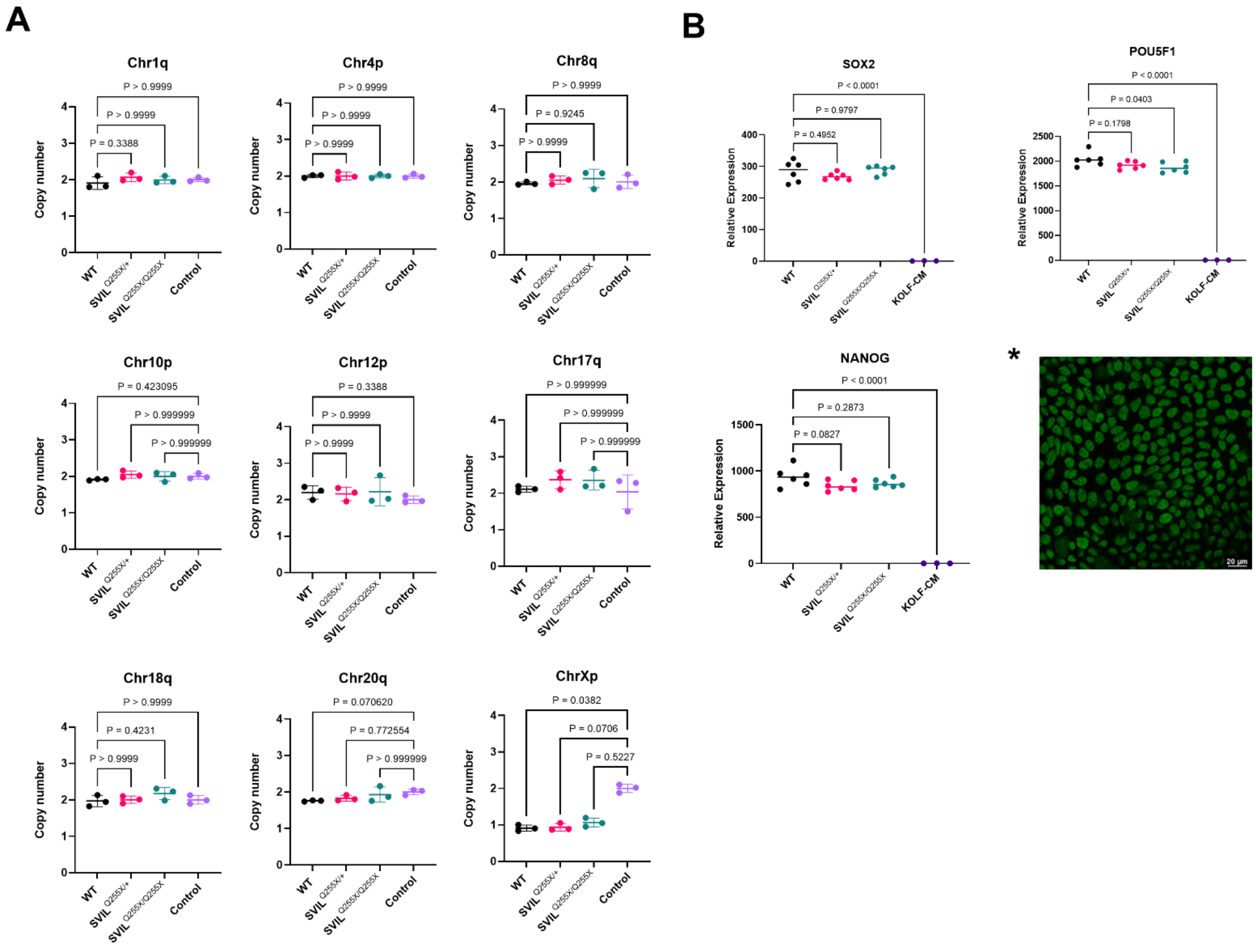


**Supplemental Figure 5: *SVIL* Q255X lines do not carry chromosomal anomalies and show normal pluripotency. (A)** Karyotyping by copy number variation using RT-qPCR. **(B)** RT-qPCR of pluripotency markers SOX2, POU5F1 and NANOG. Control: unedited iPSC KOLF2 CM. *Representative pluripotency IF for SOX nuclei staining

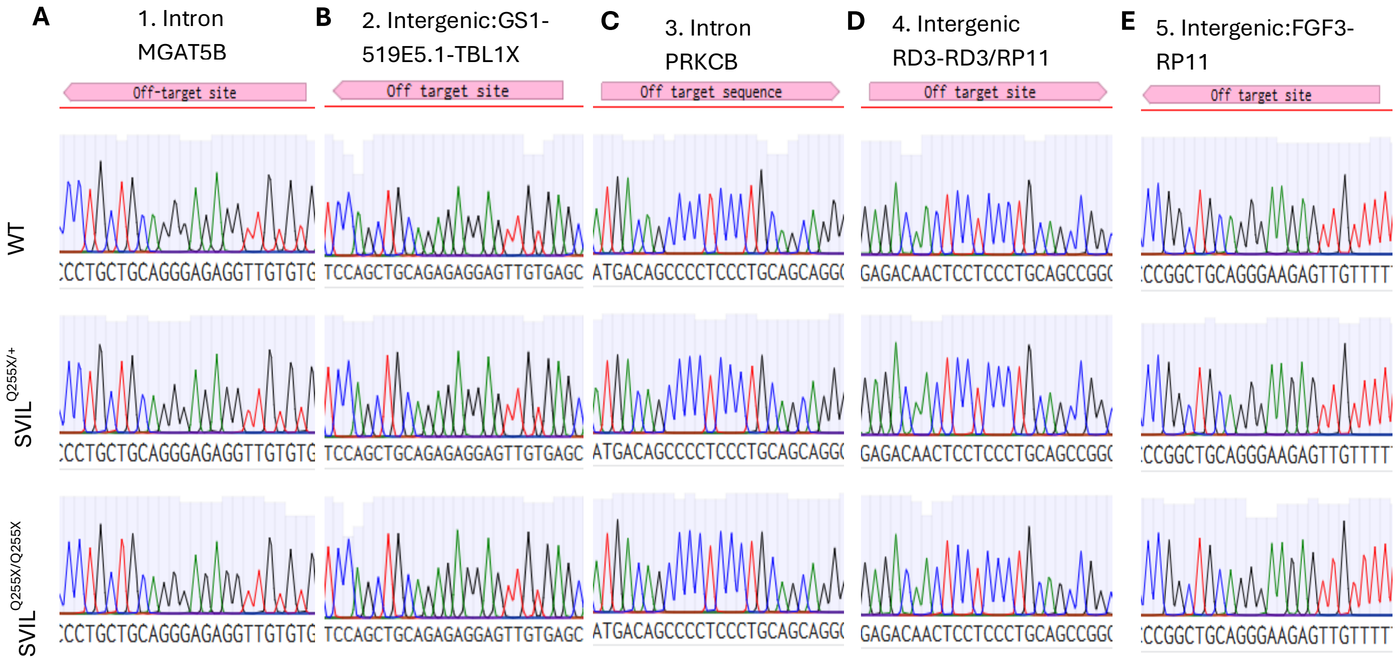


**Supplemental Figure 6: Sanger sequencing of top 5 CRISPOR predicted off-target sites confirmed no off-target editing in engineered Q255X iPSC KOLF lines**. **(A)** Intronic MGAT5B. **(B)** Intergenic between GS1-519E5.1 to TBL1X. **(C)** Intronic PRKCB. **(D)** Intergenic between RD3 and RD3/RP11. **(E)** Intergenic between FGF4 and RP11.

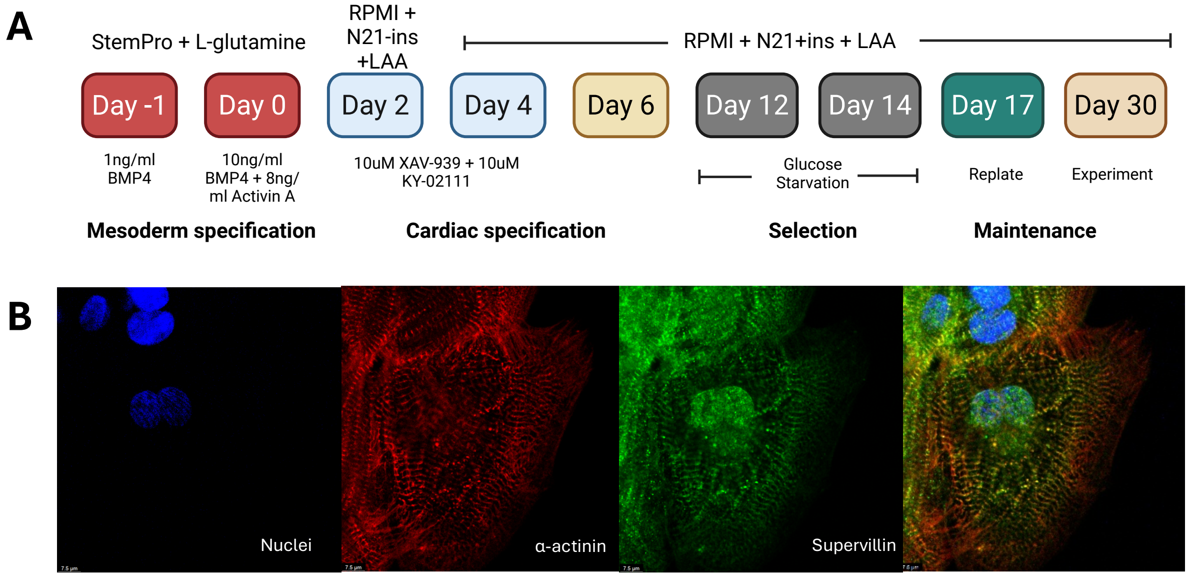


**Supplemental Figure 7: Engineered iPSCs can be successfully differentiated into cardiomyocytes. (A)** iPSC cardiomyocyte differentiation protocol using small molecule Wnt-pathway activation and inhibition. **(B)** Representative IF image showing differentiated iPSC-CM stained for nuclei using TO-PRO (blue), alpha-actinin antibody (red) and supervillin antibody (green).

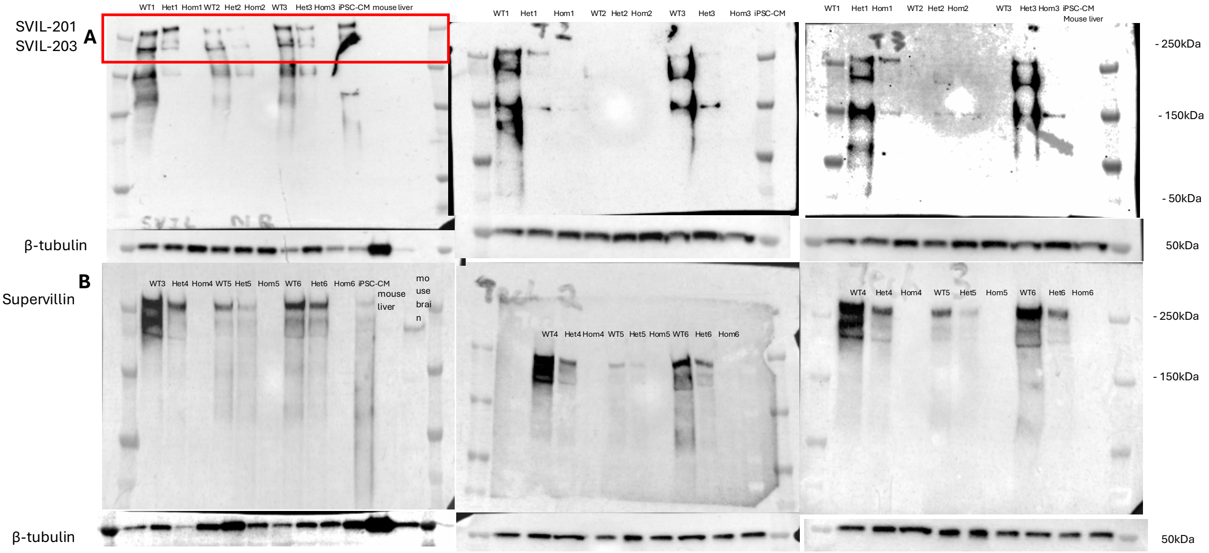


**Supplemental Figure 8: *SVIL* Q255X iPSC-CMs show protein haploinsufficiency by western blotting.** SVIL-201 (250kDa) is the skeletal and cardiac muscle-specific isoform and SVIL-203 (205kDa) is the non-muscle specific isoform. **(A)** Biological replicates 1-3 over 3 technical replicates. **(B)** Biological replicates 4-6 over 3 technical replicates. Het = $SVIL^{Q255X/+}$, Hom = $SVIL^{Q255X/Q255X}$. Positive control: unedited iPSC-CM; negative control: mouse liver; mouse brain showed alternative isoform of supervillin

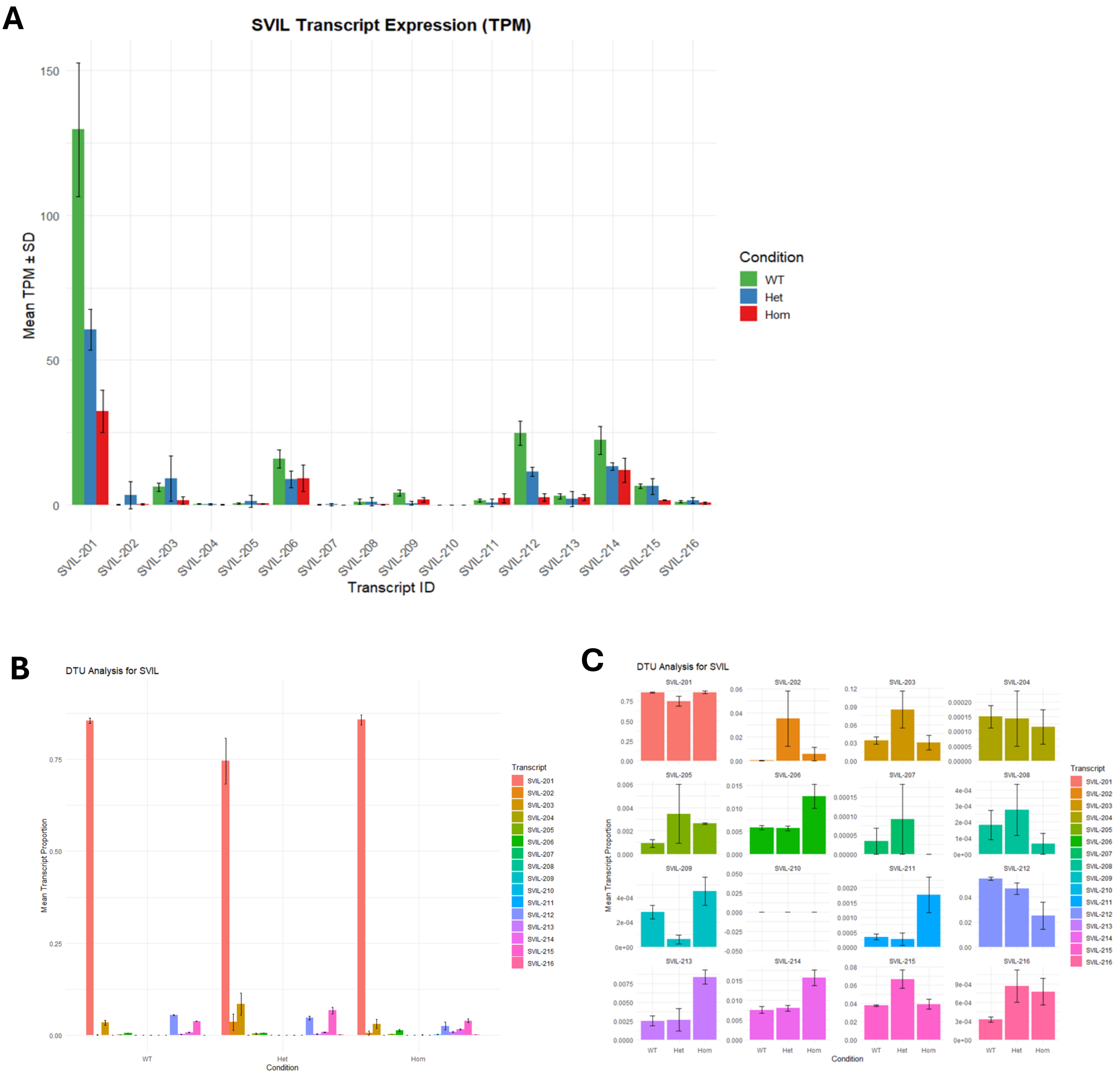


**Supplemental Figure 9: Differential transcript expression and transcript usage analysis using RNA-seq data show no evidence of isoform switching. (A)** SVIL transcript expression in transcript per million (TPM) across the different SVIL transcripts. **(B + C)** Differential Transcript Usage analysis of SVIL transcripts per genotype. Het = $SVIL^{Q255X/+}$, Hom = $SVIL^{Q255X/Q255X}$. N = 3 biological replicates for WT and $SVIL^{Q255X/Q255X}$, 4 for $SVIL^{Q255X/+}$. Bars show the mean proportion of each SVIL transcript within each genotype group with error bars indicating SEM. Kruskal-Wallis test performed with post-hoc Dunn's and BH-correction.

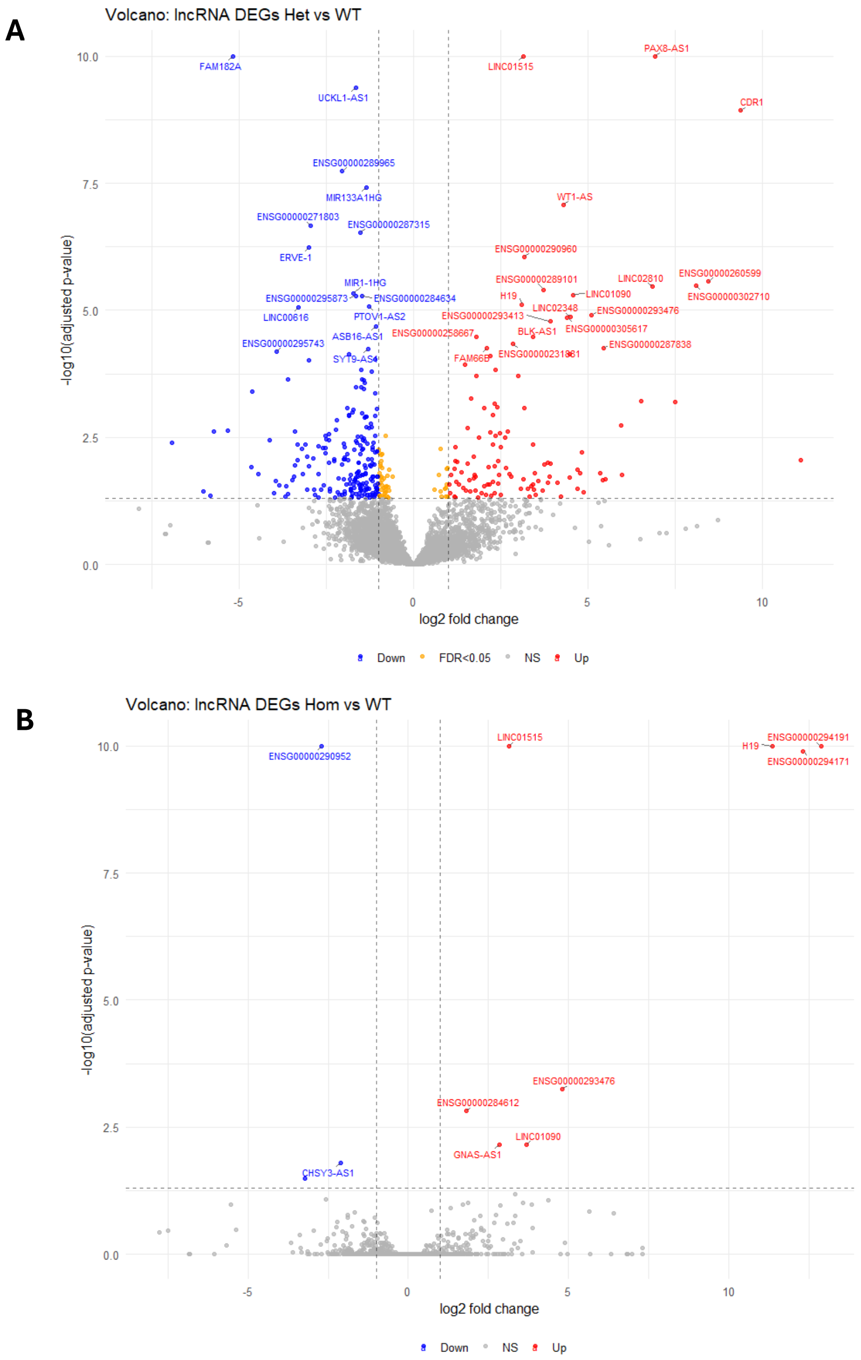


**Supplemental Figure 10: Long non-coding RNA (lncRNA) from RNA-seq shows 4 overlapping gene/proteins (H19, LINC01515, LINC01090, and GNAS-AS1) of which H19 is of interest in HCM. (A)** Volcano plot of lncRNA for *SVIL*$^{Q255X/+}$ vs WT contrast. **(B)** Volcano plot of lncRNA for *SVIL*$^{Q255X/Q255X}$ vs WT contrast. Log2 fold change (|log2FC|) cut off set at > 1, false discovery rate (FDR) cut off set at <0.05.

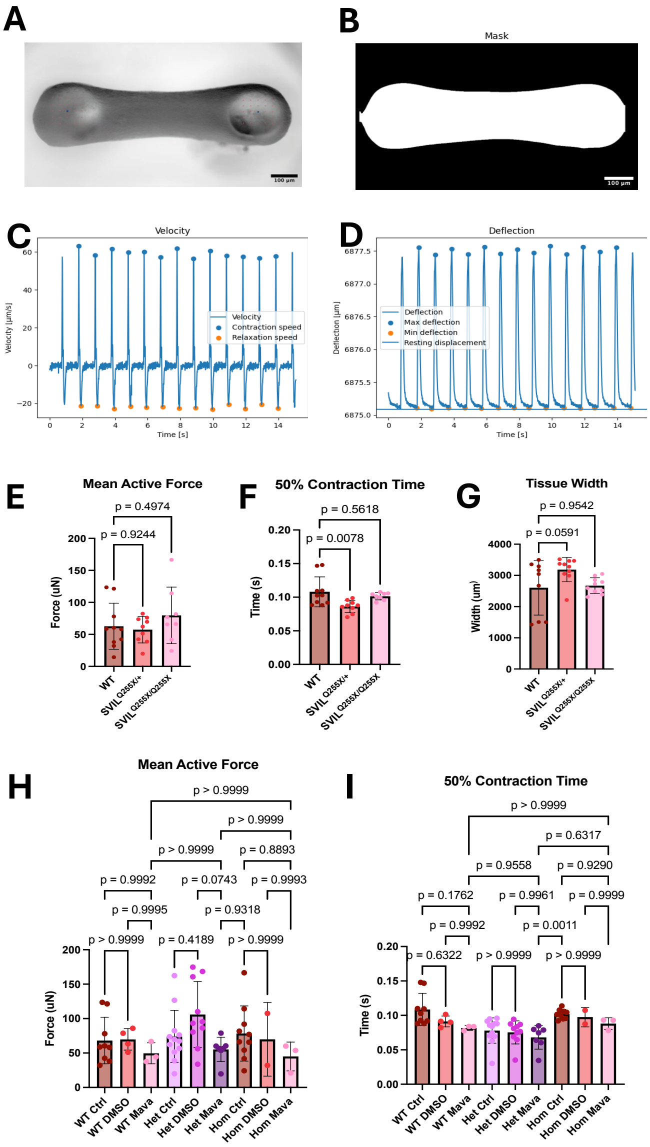


**Supplemental Figure 11: Engineered heart tissues (EHTs) using WT, SVIL$^{Q255X/+}$, and SVIL$^{Q255X/Q255X}$ iPSC-CMs. (A)** Brightfield image of EHT imaged at 2X with optical flow tracking points shown to extract functional measures of contractility. **(B)** Mask segmentation created over EHT by BeatProfiler to extract structural measures of contractility. **(C)** Example of raw calculated velocity of contractile function from BeatProfiler analysis. **(D)** Example of deflection trace calculated from optical flow tracking of the visible pillars of the EHT construct. **(E)** Mean active force of the EHTs across WT, SVIL$^{Q255X/+}$ and SVIL$^{Q255X/Q255X}$ constructs. **(F)** Time to reach 50% of contraction (50% contraction time) measured across WT, SVIL$^{Q255X/+}$ and SVIL$^{Q255X/Q255X}$ EHTs. **(G)** Measured tissue width across WT, SVIL$^{Q255X/+}$and SVIL$^{Q255X/Q255X}$ EHTs. **(H)** Effect of 30 min acute 0.3 µM Mavacamten treatment on mean active force of WT, SVIL$^{Q255X/+}$ and SVIL$^{Q255X/Q255X}$ EHTs. **(I)** The effect of 30 min acute 0.3 µM Mavacamten on time to reach 50% of contraction (50% contraction time) measured across WT, SVIL$^{Q255X/+}$ and SVIL$^{Q255X/Q255X}$ EHTs. n = 10-13 per genotype for untreated EHTs, n= 3-7 for Mavacamten treated EHTs.

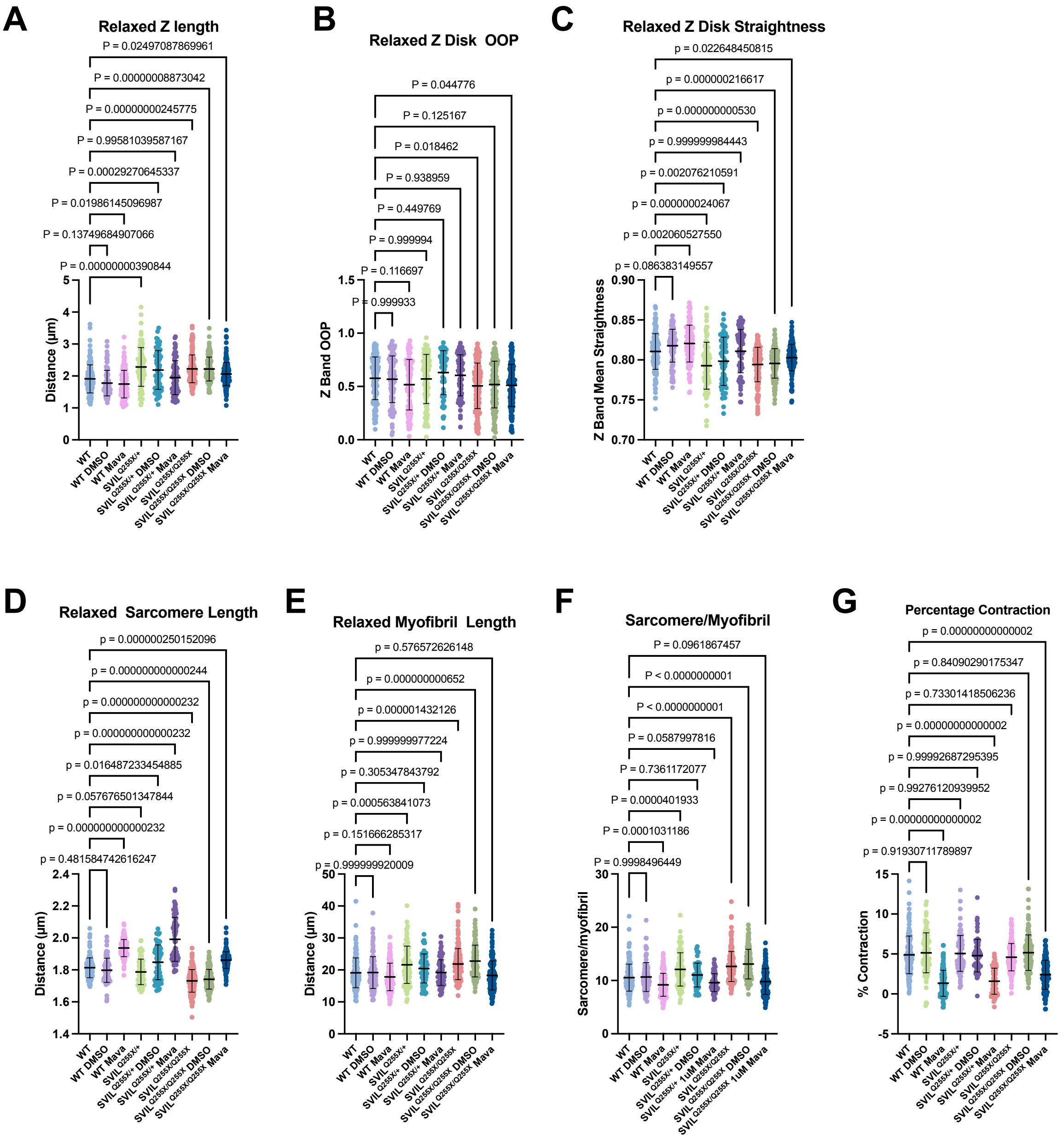


**Supplemental Figure 12: Sarcomere morphology analysis using SarcAsM for Q255X iPSC-CM lines with and without Mavacamten treatment. (A)** Relaxed sarcomere Z disc length in µm per cell. **(B)** Relaxed Z disc orientation order parameter (OOP) per cell where 1 defines fully ordered aligned sarcomeres and 0 defines randomly ordered sarcomeres. **(C)** Relaxed Z disc straightness per cell **(D)** Relaxed mean sarcomere length per cell in µm. **(E)** Relaxed myofibril length in µm per cell. **(F)** Sarcomere number per myofibril per cell. **(G)** Mean percentage contraction calculated from sarcomere length shortening per cell. Each parameter calculated from the mean of each cell sampled across WT untreated (n = 156), WT in DMSO (n = 99), WT in Mavacamten (n = 126), SVIL$^{Q255X/+}$ untreated (n = 92), SVIL$^{Q255X/+}$ in DMSO (n = 64), SVIL$^{Q255X/+}$ in Mavacamten (n = 76), SVIL$^{Q255X/Q255X}$ untreated (n = 178), SVIL$^{Q255X/Q255X}$ in DMSO (n = 133) and SVIL$^{Q255X/Q255X}$ in Mavacamten (n = 150). Cells were treated with 1 µM Mavacamten or vehicle (DMSO). Bars show mean ± standard deviation. One-way ANOVA performed with post-hoc Dunnett's multiple comparisons test.

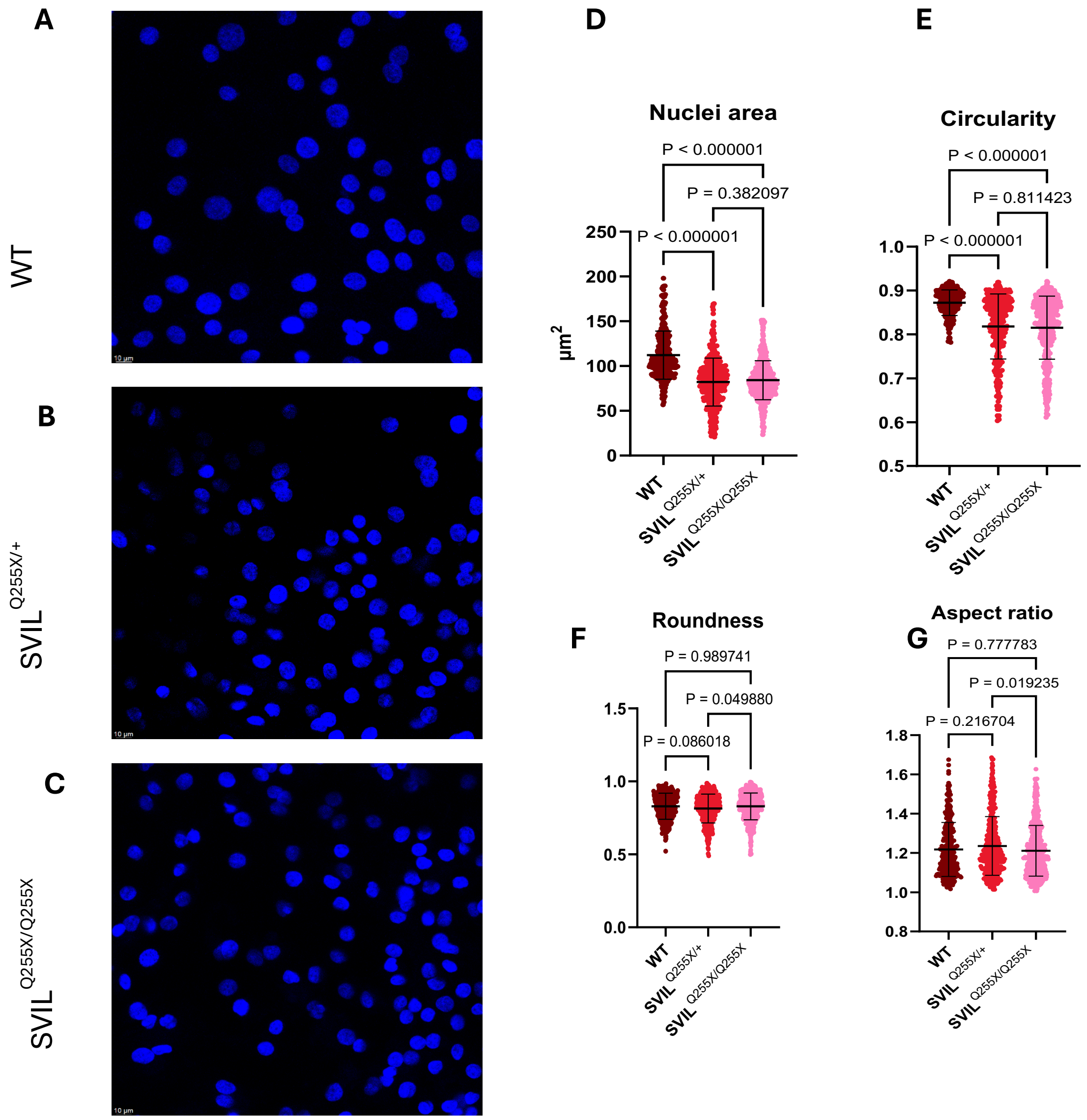


**Supplemental Figure 13: Q255X variant affects nuclear morphology of iPSC-CMs. (A)** Representative IF image of isogenic WT iPSC-CM. **(B)** Representative IF image of SVIL$^{Q255X/+}$ iPSC-CM nuclei. **(C)** Representative IF image of SVIL$^{Q255X/Q255X}$ iPSC-CM nuclei. Nuclei stained with NucBlue$^{TM}$ (Hoechst 33342). Images obtained at 63X objective. **(D)** Area per nucleus in µm$^2$ of SVIL iPSC-CM between genotype groups. WT (n = 281 nuclei), SVIL$^{Q255X/+}$ (n = 401) and SVIL$^{Q255X/Q255X}$ (n= 552). **(E)** Circularity of nuclei of SVIL iPSC-CM between genotype groups defined by 4π (area/perimeter$^2$). **(F)** Roundness of nuclei of SVIL iPSC-CMs between genotype groups defined by 4area/ π (major axis)$^2$. **(G)** Aspect ratio of nuclei of SVIL iPSC-CMs between genotype groups defined by (major axis)/(minor axis). One-way ANOVA performed with post-hoc Dunnett's multiple comparisons test.

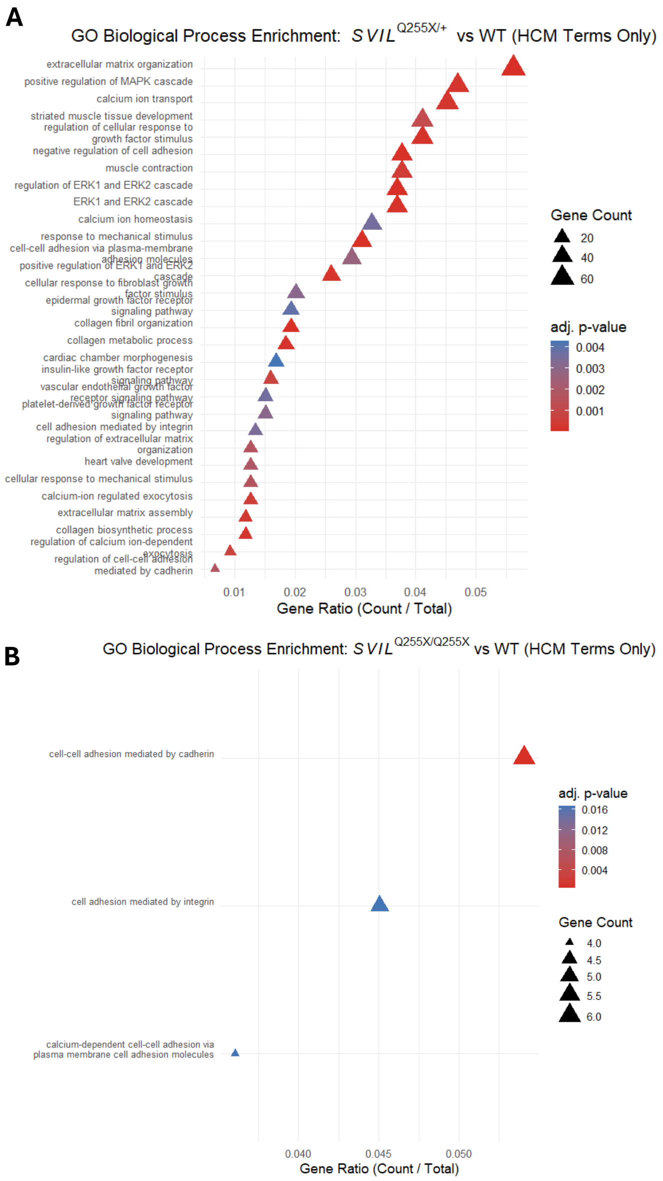


**Supplemental Figure 14: Gene Ontology Biological Process enrichment of DEGs of Q255X iPSC-CMs filtered by HCM specific keywords.** HCM keywords include: cardiomyopathy, cardiomyocyte, cardiac muscle, cardiac, heart, ventricle, atrial, myocardium, cardiac development, heart development, sarcomere, myofibril, myofilament, actin, myosin, contractility, muscle contraction, actomyosin, Z disc, cytoskeletal organization, striated muscle, calcium, calcium ion, calcium homeostasis, calcium signalling, excitation-contraction, ryanodine, troponin, SERCA, calsequestrin, fibrosis, extracellular matrix, collagen, matrix organization, matrisome, fibroblast, TGF-beta, ECM, collagen fibril, fibril organization, costamere, integrin, cell adhesion, focal adhesion, dystrophin, vinculin, talin, cytoskeleton-membrane, MAPK, ERK, p38, JNK, MAP kinase, signal transduction, growth factor, receptor signalling, stress-activated MAPK, mechanotransduction, hypertrophy, response to stress, cardiac hypertrophy, load-induced, stretch, mechanical stimulus, mechanical stress. **(A)** Enriched GO-BPs for DEGs between the *SVIL*$^{Q255X/+}$ vs WT contrast filtered by HCM-specific terms. **(B)** Enriched GO-BPs for DEGs between the *SVIL*$^{Q255X/Q255X}$ vs WT contrast filtered by HCM-specific terms. GO-BP over-representation analysis was performed on DEGs using clusterProfiler with DEGs defined by padj <0.05 and |log2 FC| >1

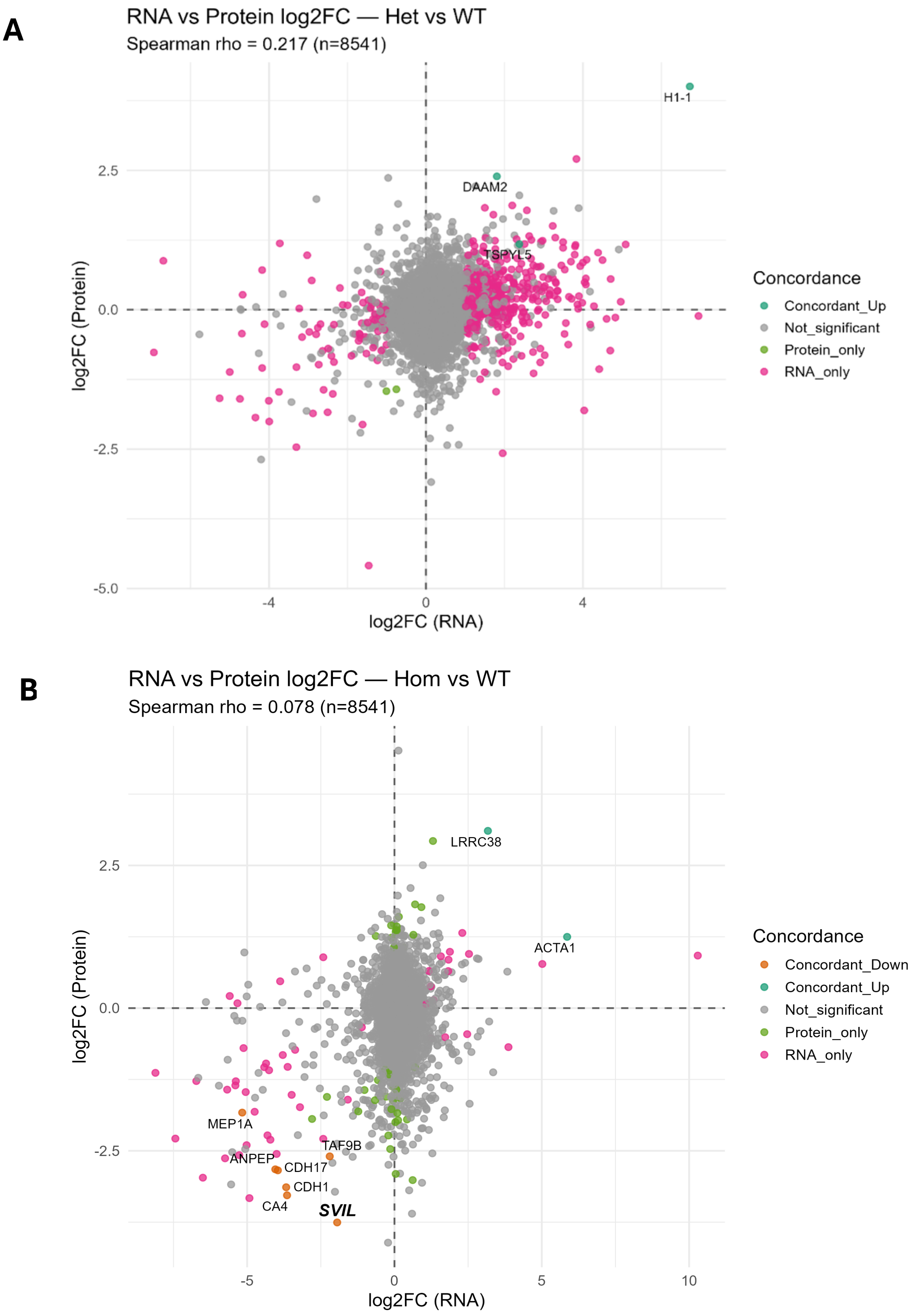


**Supplemental Figure 15: Integration analysis of omics data combining RNA-seq with proteomic data showing concordant genes/proteins. (A)** Integrated RNA vs Protein data for *SVIL*$^{Q255X/+}$ vs WT contrast showing concordant upregulated H1-1, DAAM2 and TSPYL5. **(B)** Integrated RNA vs Protein data for *SVIL*$^{Q255X/Q255X}$ vs WT contrast showing concordant upregulated LRRC38 and ACTA1 and downregulated MEP1A, TAF9B, ANPEP, CDH17, CDH1, CA4, and SVIL. No discordant gene/protein was found.

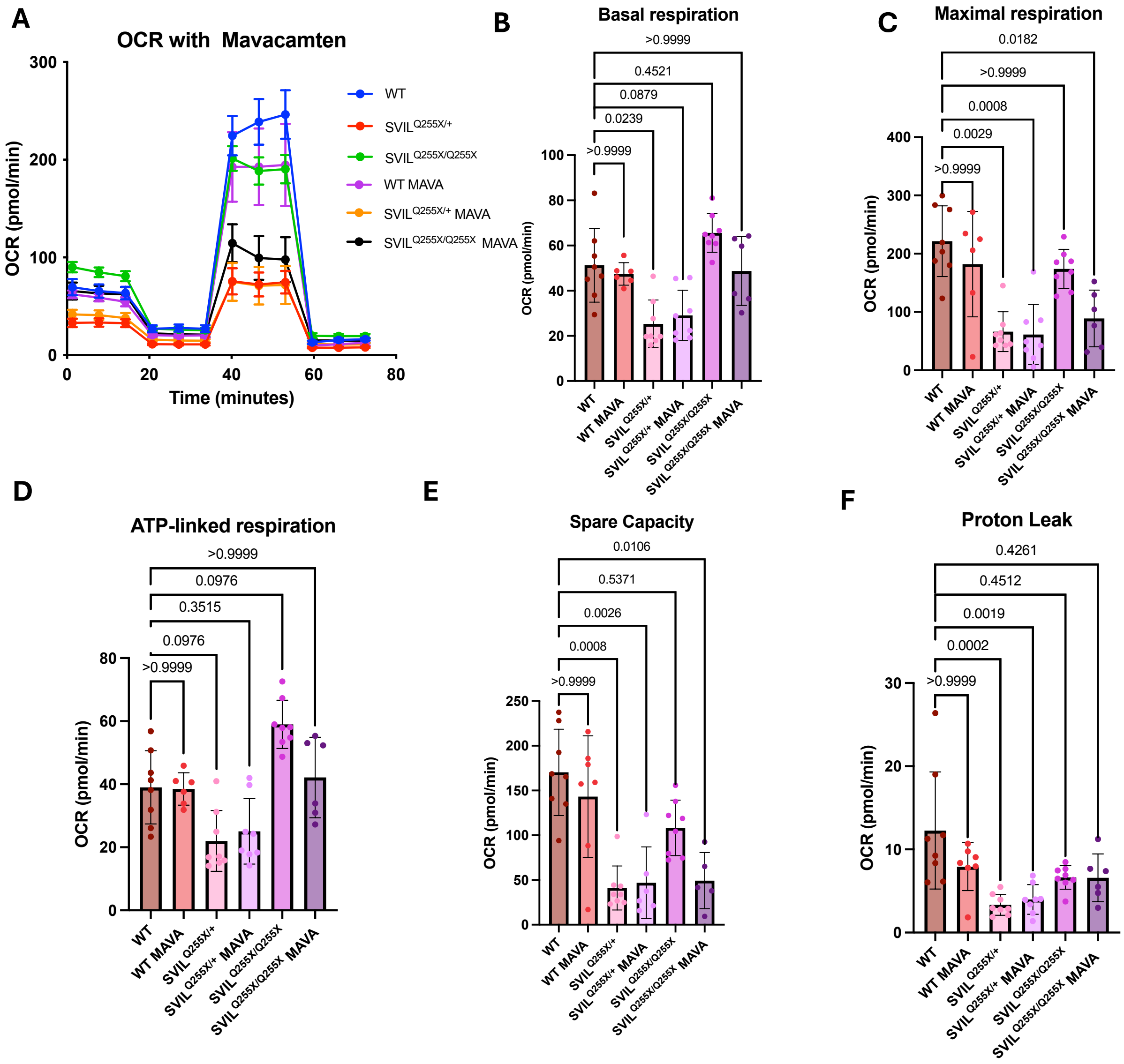


**Supplemental Figure 16: Oxygen consumption rate (OCR) with Mavacamten treatment for Q255X iPSC-CMs. (A)** Averaged and normalized OCR measured across untreated and treated WT, SVIL$^{Q255X/+}$, and SVIL$^{Q255X/Q255X}$ iPSC-CMs. Cells were treated with 1 μM Mavacamten. N = 8 technical replicates for each genotype group with trends confirmed over 2 biological replicates. Each point is plot as mean ± standard deviation. Outliers were removed using the Robust regression and Outlier removal (ROUT) method with Q = 1%. **(B)** Dot plot of basal respiration per well sampled across untreated and treated WT, SVIL$^{Q255X/+}$, and SVIL$^{Q255X/Q255X}$ iPSC-CMs. Bars showing mean ± standard deviation. **(C)** Dot plot of maximal respiration per well sampled across untreated and treated WT, SVIL$^{Q255X/+}$, and SVIL$^{Q255X/Q255X}$ iPSC-CMs. Bars showing mean ± standard deviation. **(D)** Dot plot of ATP-linked respiration per well sampled across untreated and treated WT, SVIL$^{Q255X/+}$, and SVIL$^{Q255X/Q255X}$ iPSC-CMs. Bars showing mean ± standard deviation. **(E)** Dot plot of proton leak per well sampled across WT, SVIL$^{Q255X/+}$, and SVIL$^{Q255X/Q255X}$ iPSC-CMs. Bars showing mean ± standard deviation. **(F)** Dot plot of maximal respiration per well sampled across WT, SVIL$^{Q255X/+}$, and SVIL$^{Q255X/Q255X}$ iPSC-CMs. Bars showing mean ± standard deviation.

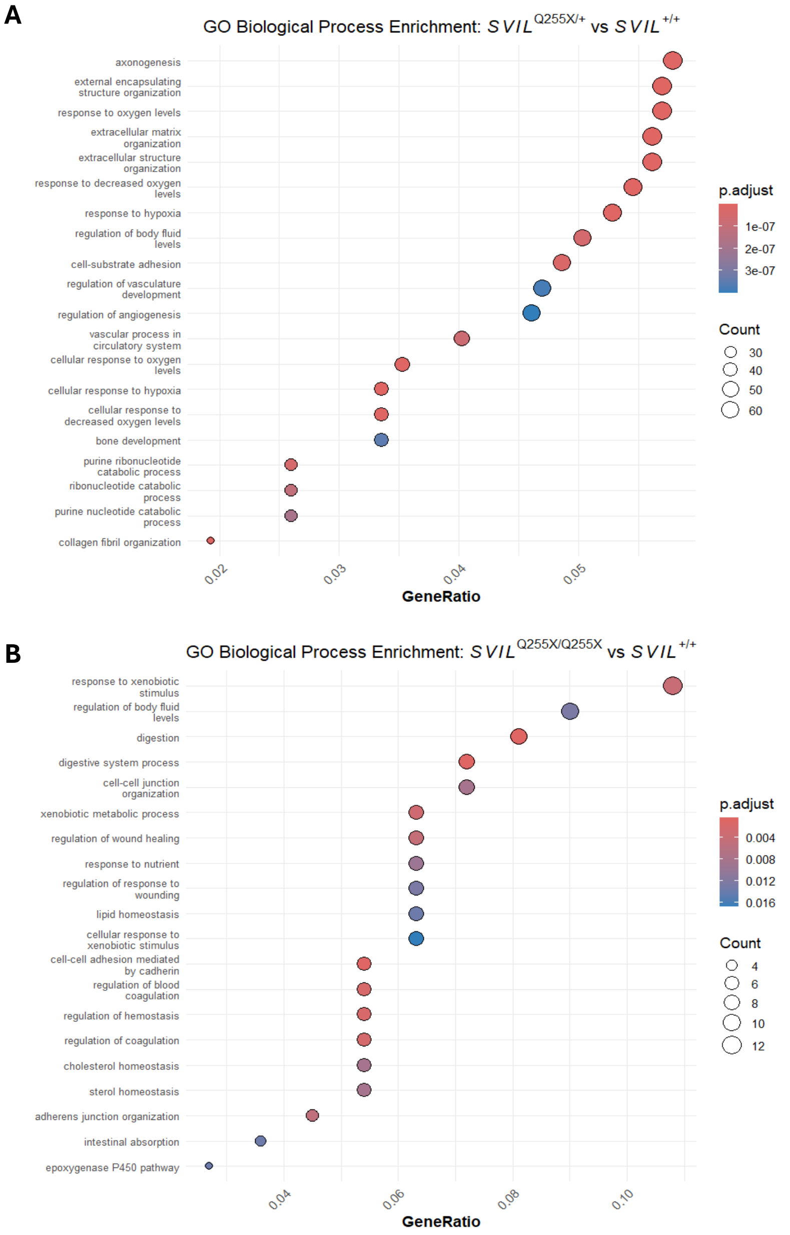


**Supplemental Figure 17: Unbiased RNA-seq pathway enrichment using Gene Ontology Biological Process (GO-BP) show hypoxia and ECM related pathways in *SVIL*$^{Q255X/+}$ vs WT which is not present *SVIL*$^{Q255X/Q255X}$ vs WT. (A)** GO-BP pathways for *SVIL*$^{Q255X/+}$ vs WT contrast showing 6 out of 20 most enriched pathways relating to hypoxia, 5 relating to ECM, and 3 relating to angiogenesis and vasculature. **(B)** GO-BP pathways for *SVIL*$^{Q255X/Q255X}$ vs WT contrast do not show HCM relevant pathways
